# Pioneer factor ETV2 safeguards endothelial cell specification by recruiting the repressor REST to restrict alternative lineage commitment

**DOI:** 10.1101/2024.05.28.595971

**Authors:** Danyang Chen, Xiaonuo Fan, Kai Wang, Liyan Gong, Juan M. Melero-Martin, William T. Pu

## Abstract

Mechanisms of cell fate specification remain a central question for developmental biology and regenerative medicine. The pioneer factor ETV2 is a master regulator for the endothelial cell (EC) lineage specification. Here, we studied mechanisms of ETV2-driven fate specification using a highly efficient system in which ETV2 directs human induced pluripotent stem cell-derived mesodermal progenitors to form ECs over two days. By applying CUT&RUN, single-cell RNA-sequencing (scRNA-seq) and single-cell assay for transposase-accessible chromatin sequencing (scATAC-seq) analyses, we characterized the transcriptomic profiles, chromatin landscapes, dynamic cis-regulatory elements (CREs), and molecular features of EC cell differentiation mediated by ETV2. This defined the scope of ETV2 pioneering activity and identified its direct downstream target genes. Induced ETV2 expression both directed specification of endothelial progenitors and suppressed acquisition of alternative fates. Functional screening and candidate validation revealed cofactors essential for efficient EC specification, including the transcriptional activator GABPA. Surprisingly, the transcriptional repressor REST was also necessary for efficient EC specification. ETV2 recruited REST to occupy and repress non-EC lineage genes. Collectively, our study provides an unparalleled molecular analysis of EC specification at single-cell resolution and identifies the important role of pioneer factors to recruit repressors that suppress commitment to alternative lineages.

## Introduction

Diverse cell lineages are formed from a single multipotent progenitor during mammalian development. Elucidating the mechanisms responsible for cell lineage specification and differentiation is a fundamental goal of developmental biology, and a better understanding of these mechanisms will have direct applications in regenerative medicine. We previously reported that expression of ETV2, a master regulator of endothelial cell (EC) specification, directs mesoderm progenitor cells to rapidly and efficiently differentiate into ECs^1^. Here, we further optimize this powerful lineage specification model system and deploy it to dissect mechanisms by which a master transcriptional regulator programs progenitor cells to specify progeny lineages.

DNA occupancy by nucleosomes is a key epigenetic barrier that constrains cells to a restricted set of transcriptional states and lineages^2^. Pioneer transcription factors (TFs) can occupy binding sites within nucleosomal DNA and make it accessible for binding by non-pioneer factors^3^. This property makes pioneer TFs central to lineage specification, as their expression allows cells to surmount genome accessibility constraints to differentiate into additional cell types^4^. For instance, the prototypical pioneer TFs FOXA1/2 are required in early endodermal cells for hepatocyte specification^5^. FOXA1/2 binds distal enhancers and activates the expression of genes that dictate hepatocyte fate, such as the non-pioneer TF HNF4a^6^.

Analogous to the FOXA1/2 for the liver, ETV2 is a pioneer TF that is critical for EC specification^7–10^. This ETS-family TF is transiently expressed in mesodermal EC progenitors from embryonic day (E) 7.75-E9.5 of mouse development. Mice lacking ETV2 fail to develop blood or vasculature^11,12^, and induced pluripotent stem cells (iPSCs) lacking ETV2 do not form ECs^1^. Forced expression of ETV2 directs iPSCs, fibroblasts, and mesoderm progenitor cells (MPCs) to differentiate into ECs^1,13–15^. Among ETV2’s identified target genes are hallmark EC genes *Tie2*, *Lmo2*, and *Fli1*^8,12,16^. Despite this large body of data, the mechanisms by which ETV2 directs mesodermal progenitors to differentiate into ECs remain unclear.

Here, we developed a highly efficient and rapid iPSC-based system to study ETV2-driven EC specification and differentiation from MPCs. We used this system with single-cell approaches to determine ETV2’s impact on MPC differentiation trajectories. We identified enhancers and genes regulated by ETV2 pioneering activity and uncovered TFs that collaborate with ETV2 to direct EC specification. We discovered that ETV2 recruits REST, a transcriptional repressor, to silence non-EC lineage genes, enhancing EC specification by blocking alternative fates. Collectively, our study provides molecular insights into the mechanisms by which ETV2 drives endothelial specification.

## Results

### Highly efficient and rapid ETV2-directed differentiation of iPSC-derived MPCs to ECs

We first established a system to rapidly, efficiently, and reproducibly induce iPSC-derived MPCs to differentiate into ECs. We previously showed that modified mRNA-mediated expression of ETV2 in iPSC-derived MPCs efficiently directs their differentiation to iPSC-derived ECs^1^. To increase the reproducibility and flexibility of programmed ETV2 expression, we engineered a stable clonal iPSC line, TRE3G-ETV2 iPSC, that inducibly expresses ETV2 upon treatment with doxycycline (Fig. 1A). Treatment of iPSCs with Wnt activator CHIR99021 for two days induced formation of MPCs. On Day 2 (D2), CHIR99021 was removed, and cells were cultured for two additional days in the presence of angiogenic growth factors (VEGFA, FGF2, and EGF) and the TGFβ inhibitor SB413542. In the absence of ETV2 induction (Ctrl), this protocol yielded ∼30% CDH5 (VE-cadherin) and PECAM1 double positive iPSC-ECs, as measured by fluorescence-activated cell sorting (FACS; Fig. 1A-C). Treatment with Dox from D2-D3 most efficiently induced iPSC-ECs (94% CDH5^+^ PECAM1^+^), whereas shifting Dox treatment to D3-D4, extending the duration from D2-D4, or extending CHIR99021 to D3 followed by Dox treatment were less efficient (Fig. 1A-C and Extended Data Fig. 1A-C). We examined the effect of varying the dose of Dox, and found a steeply non-linear response consistent with the dose-response of reverse tet-activator systems^17^ (Extended Data Fig. 1D,E). RT-qPCR verified that *ETV2* mRNA levels were significantly upregulated by Dox treatment for 24 hours (Extended Data Fig. 1F). These results yielded an optimized four-day protocol in which ETV2 induction by 1 µg/ml Dox from D2-D3 resulted in highly efficient MPC-to-EC differentiation by D4.

**Fig. 1.**
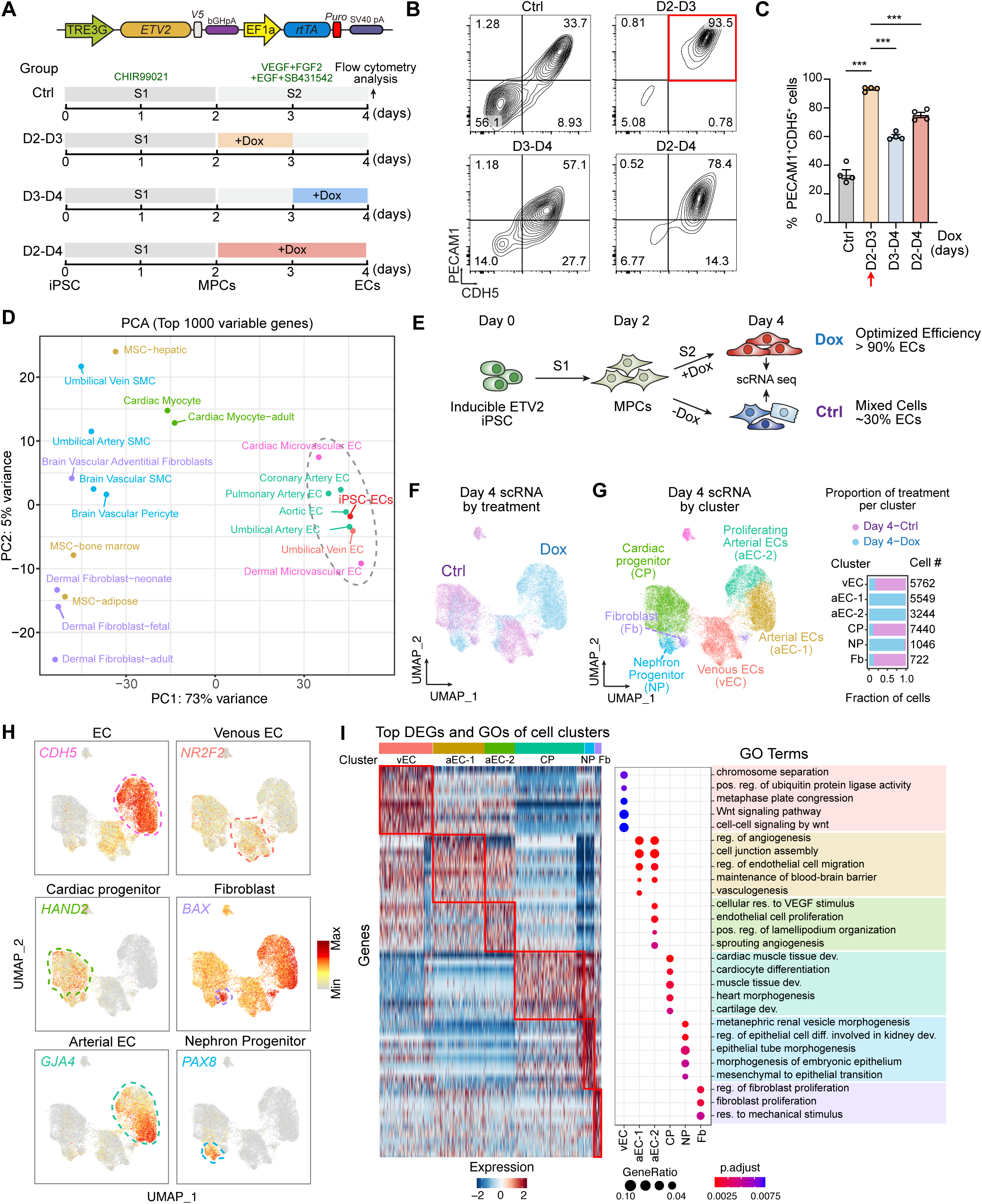
Highly efficient and rapid EC differentiation from mesodermal progenitors by ETV2 induction. (**A**) Overview of the TRE3G-ETV2-V5 system (upper panel) and timeline of 4 day EC differentiation protocol showing timing of Dox addition (1 μg/ml). Cells were treated without Dox (Ctrl), or treated with Dox for indicated length of time. **(B,C)** Flow cytometry analysis (B) and quantification (C) for PECAM1 and CDH5 expression at day 4 of differentiation. One-way ANOVA, Tukey’s multiple comparison test. n = 4. **(D)** PCA plot based on the top 1,000 most variable genes across iPSC-ECs and indicated human cell types (from GSE138734). **(E)** Timeline of the optimized 4 day EC differentiation protocol. Cell samples were collected from Ctrl and Dox groups on day 4 for scRNA seq. **(F)** scRNA-seq gene expression projected onto a UMAP space and labeled by group (Ctrl, Dox). **(G)** scRNA-seq labeled by inferred cell types. Bar plots show the proportion of cells from each treatment across the clusters. **(H)** Feature plots of selected markers of each cluster. **(I)** Heatmap of the top DEGs in each cluster and their most significantly enriched GO terms. MPCs, mesoderm progenitor cells; ECs, endothelial cell; reg, regulation; dev, development; pos, positive. ***, P<0.001.

We characterized the iPSC-ECs generated using this optimized protocol by positively selecting cells on D4 using anti-PECAM1-conjugated magnetic beads. Immunostaining for CDH5 and PECAM1 showed that these EC markers had the expected cell surface expression (Extended Data Fig. 2A). The cells aligned in response to flow, indicating that they can sense and respond to fluid shear stress (Extended Data Fig. 2B). The iPSC-ECs also exhibited expected EC phenotypes: they migrated in the wound scratch assay (Extended Data Fig. 2C), produced nitric oxide (Extended Data Fig. 2D), took up acetylated low-density lipoprotein (AcLDL; Extended Data Fig. 2E), and upregulated *ICAM1*, *VCAM1*, and *E-selectin (SELE)* in response to the inflammatory cytokine tumor necrosis factor-alpha (Extended Data Fig. 2F).

We then compared the transcriptional similarity of the generated iPSC-ECs to primary cells isolated from human tissues, including diverse types of ECs, smooth muscle cells, fibroblasts, and cardiomyocytes, using RNA-seq data^18^. Principal component analysis (PCA) based upon the top 1000 variable genes revealed that the iPSC-ECs grouped with ECs and away from non-ECs (Fig. 1D), suggesting an overall transcriptional similarity with primary ECs. The iPSC-ECs had the greatest transcriptional similarity with arterial ECs (aECs). Collectively, these studies confirmed that ETV2-directed, iPSC-derived ECs possess functional properties of ECs in vitro and are transcriptionally most similar to aECs.

### scRNA-seq comparison of iPSC-EC differentiation with and without ETV2 overexpression

To better characterize the products of control and Dox differentiation, we acquired high-quality single-cell transcriptomes from 25,502 control (Ctrl: no Dox) and Dox cells at differentiation D4 (Fig. 1E and Supplementary Table 1). We used uniform manifold approximation and projection (UMAP) to visualize the results. The majority of Dox and Ctrl cells at D4 were transcriptionally distinct (Fig. 1F). Graph-based clustering identified six cell clusters (Fig. 1G). Ctrl cells predominantly belonged to three clusters. One of these was a venous EC (vEC) cluster, as it expressed pan-EC markers *CDH5*, *PECAM1*, and *ESAM* and vEC markers *NR2F2*, *APLNR*, and *EPHB4* (Fig. 1G,H). The remaining Ctrl cells did not express EC markers, and rather expressed markers of cardiac progenitors (*HAND2*) or fibroblasts (*BAX*). In contrast, most Dox cells contributed to two large, related clusters (aEC-1 and aEC-2) that highly expressed pan-EC markers and aEC genes^19^ such as *GJA4*, *DLL4*, and *EFNB2* (Fig. 1G,H). A small fraction of Dox cells did not express EC markers. Most of these cells expressed genes such as *PAX8* and *PAX2*, characteristic of nephron progenitors (NP; Fig. 1G,H). These results show that Ctrl and Dox cells yielded distinct and largely non-overlapping cell types, with Dox cells nearly exclusively forming aECs and NPs, and Ctrl cells predominantly forming vECs, fibroblasts, and cardiac progenitors (Fig. 1G).

Gene ontology (GO) enrichment analysis of the marker genes of each cluster further revealed their biological function and cell identity (Fig. 1I). For example, the aEC clusters from Dox cells were enriched for functional terms related to the regulation of angiogenesis, cell junction assembly, and regulation of EC migration whereas the cardiac progenitor cluster, largely from Ctrl cells, was enriched for cardiac terms such as cardiac muscle tissue development and heart morphogenesis.

Taken together, these results confirm that ETV2 efficiently differentiates MPCs into iPSC-ECs.

### Single-cell analysis of iPSC-EC differentiation dynamics

To assess chromatin and gene expression dynamics during EC specification, we performed separate scATAC-seq and scRNA-seq at days 2 (D2), 3 (D3), 3.5 (D3.5), and 4 (D4) from the Ctrl and Dox differentiation conditions. After stringent filtering and quality control, we obtained 59,785 high-quality scATAC-seq cells and 105,563 high-quality scRNA-seq cells with excellent overlap between biological replicates (Extended Data Fig. 3A-C and Supplementary Table 1). These cells exhibited a high signal-to-noise ratio at transcription start site (TSS) and a canonical fragment-size distribution (Extended Data Fig. 3B). For scATAC-seq and scRNA-seq datasets, we performed integrative analysis of Ctrl and Dox cells at the four time points (Fig. 2A). The D2 cells, representing mesoderm progenitor cells, from Ctrl and Dox time courses grouped together as expected, suggesting that batch effects were insignificant. By contrast, after dox treatment, Dox cells at each subsequent time point largely clustered in a different UMAP space than Ctrl (Fig. 2A). This indicates that ETV2 guides EC specification by reshaping the genomic accessibility landscape and gene expression.

**Fig. 2.**
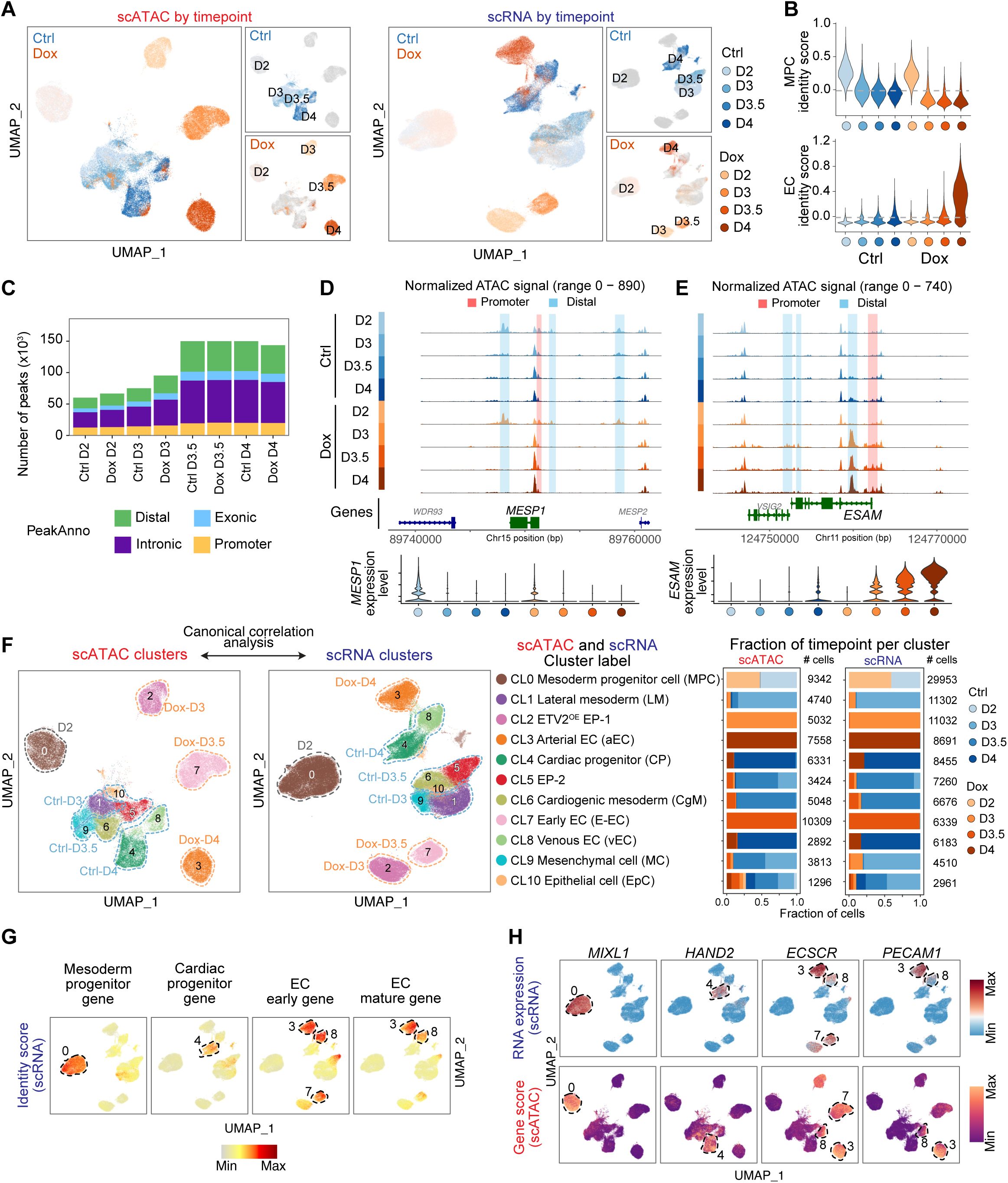
Integrated single-cell transcriptomic and epigenomic analysis of EC differentiation. (**A**) Time course of scATAC-seq and scRNA-seq data from D2, D3, D3.5 and D4 projected onto UMAP space showing days of differentiation of both Ctrl and Dox cells. Cells are colored by sample collection time. **(B)** Cell identity scores of mesoderm and EC gene sets within each timepoint. **(C)** The number of peaks for each timepoint. Bars are colored by peak location relative to genomic annotations. **(D,E)** Genome tracks of time point-resolved aggregate scATAC-seq data around the *MESP1* (D) and *ESAM* (E) gene loci. Expression of genes represented in bottom panel from scRNA-seq data. **(F)** Left panel, UMAP from scATAC-seq and scRNA-seq colored by identified cell clusters. Middle panel indicates biological classification of the scRNA-seq and inferred scATAC-seq clusters, based on known markers. Right panel, contribution of each sample to each scATAC-seq and scRNA-seq cluster. **(G)** Feature plots showing gene set identity scores. Each cell’s value is shown, mapped onto the scRNAseq UMAP plot. **(H)** Feature plots showing gene expression (top row) and gene accessibility scores (bottom row) of cell identity markers. Each cell’s value is shown, mapped onto either the scRNAseq UMAP plot (top row) or the scATACseq UMAP plot (bottom row). MPC, mesoderm progenitor cell; ETV2^OE^, ETV2 overexpression; EP, endothelial progenitor.

To investigate the differentiation dynamics, we analyzed gene expression at each time point (Extended Data Fig. 3D). Activation of WNT by CHIR99021 differentiated iPSCs into *MIXL1*^+^ and *DLL3*^+^ mesoderm at D2 while suppressing ectoderm and endoderm formation (Extended Data Fig. 3D). Consistent with this analysis, composite cell identity scores^20^ likewise demonstrated loss of MPC identity by D3 in both Ctrl and Dox (Fig. 2B). EC identity progressively increased thereafter. Early EC markers including *KLF2*, *KDR*, *ECSCR,* and *HHEX* were acquired by Dox cells by D3. Subsequently, Dox cells expressed mature EC markers including *PECAM1*, *FLT1*, *ESAM* and *JAM3* at D3.5 and D4. At D4, Dox cells expressed aEC markers (Extended Data Fig. 3D). The shift towards EC identity was only weakly apparent in Ctrl cells. Dox cells had a strong EC identity signal at D4, and this was substantially weaker in Ctrl cells (Fig. 2B). These pseudobulk analyses of the scRNA-seq data were consistent with the FACS-based measurements of EC differentiation efficiency and the D4 endpoint scRNA-seq evaluation of Ctrl and Dox cells.

Next, we performed an initial analysis of the scATAC-seq data. As expected for high-quality ATAC-seq data^21^, open chromatin regions were mostly located at intronic and intergenic regions, although promoters were over-represented compared to their small contribution to the genome (Fig. 2C). Open chromatin region number increased between D2 and D3.5, potentially due to cell state transition. Review of the scATAC-seq signal at *MESP1*, *ESAM*, and *GJA4* provided further confidence in the data (Fig. 2D,E and Extended Data Fig. 3E). *MESP1*, a master TF of mesoderm progenitors, was expressed at D2 and thereafter downregulated. Consistent with this expression pattern, D2 accessible distal regions flanking *MESP1* lost accessibility at later time points, suggesting enhancer inactivation (Fig. 2D). *ESAM*, a pan-EC gene, was progressively upregulated between D3-D4, and this upregulation was markedly stronger in Dox compared to Ctrl. In keeping with this expression profile, an intronic region of *ESAM* was closed at D2 and became highly accessible at later time points in Dox cells, suggestive of an EC enhancer. This region was only weakly accessible in D3.5 and D4 Ctrl data (Fig. 2E). *GJA4*, an aEC marker gene, was only expressed in Dox samples at D4, corresponding to increased accessibility of the promoter and a distal enhancer at D3.5 and D4 in Dox cells (Extended Data Fig. 3E).

Unsupervised clustering of the combined scRNA-seq datasets defined 11 distinct clusters (CL0-CL10) (Supplementary Table 2). To integrate the chromatin accessibility and transcriptomic profiles into a unified cell-type-resolved regulatory atlas, we performed canonical correlation analysis^22^ to match scATAC-seq cells with their nearest neighbor cells in the annotated scRNA-seq data. Most cells in scATAC-seq clusters mapped to a single corresponding scRNA-seq cluster (Extended Data Fig. 3F), and corresponding scRNA-seq and scATAC-seq clusters comprised concordant proportions across cell groups and time points (Fig. 2F, Fraction of timepoint per cluster). Dox cells were distinct from Ctrl cells after D2, and Dox cells were represented by a single dominant cluster at D3, D3.5 and D4, indicating the relative homogeneity of Dox cells at each time point: D3: CL2, ETV2 overexpression endothelial progenitor (EP)-1 (hereafter refer as ETV2^OE^ EP-1); D3.5: CL7, early EC; and D4: CL3, aECs. In contrast, the Ctrl cells had higher cellular heterogeneity: D3: CL1, lateral mesoderm, and CL9, mesenchymal cell; D3.5: CL5, EP-2, and CL6, cardiogenic mesoderm; and D4: CL4, cardiac progenitor, and CL8, vEC (Fig. 2F).

We further validated the assigned cluster identities. Consistent with their assigned labels, identity scores calculated for mesoderm progenitor, cardiac progenitor, and EC gene sets selectively highlighted CL0 (D2 cells), CL4, and Clusters 3, 7, 8, respectively (Fig. 2G). Known signature genes for each cell population, including *MIXL1* for mesodermal progenitors, *HAND2* for cardiac progenitors, *ECSCR* for early ECs identity, and *PECAM1* for mature ECs identity, were expressed most highly in the respective clusters (Fig. 2H, top row). Based on the correlation between chromatin accessibility and gene expression, we identified 77,680 significant peak-to-gene linkages including promoter and putative distal enhancers across differentiation (Extended Data Fig. 3G and Supplementary Table 3). The scATAC-seq gene accessibility scores of these signature marker genes were concordant with marker expression (Fig. 2H, bottom row).

Collectively, these results revealed transcriptomic and epigenomic signatures of iPSC-ECs, demonstrating that dual-omics mapping broadly captures major iPSC-EC cell types and the clustering outcome using scRNA-seq and scATAC-seq are concordant.

### Endogenous and ETV2-directed EC differentiation trajectories

To investigate the cell differentiation dynamics of MPCs (CL0) with or without ETV2 overexpression, we reconstructed cell trajectories in Ctrl and Dox groups (Fig. 3A). Under Dox treatment, MPCs followed a single major trajectory to differentiate into ETV2^OE^ EP-1 (CL2; D3), then into early ECs (CL7; D3.5), and finally into aECs (CL3; D4; Fig. 3B). Under Ctrl conditions, MPCs primarily differentiated into lateral mesoderm (CL1; D3; Fig. 3C). Trajectories then bifurcated, with one terminating on cardiac progenitors (CL4; D4), and the other on vECs (CL8; D4). The trajectory analysis highlighted the dramatic effect of forced ETV2 expression in MPCs: early forced ETV2 expression in the Dox group (Fig. 3D) drove differentiation along a distinct trajectory, forming EC progenitors that ultimately yielded aECs, compared to Ctrl, in which MPCs differentiated into additional mesoderm progenitors, which then formed either cardiac progenitors or vECs. Based on the relative proportion of Ctrl cells in vECs versus cardiac progenitors clusters at D4, most cells follow the cardiac rather than the EC trajectory, and as a result EC differentiation is inefficient in Ctrl cells (Fig. 3C).

**Fig. 3.**
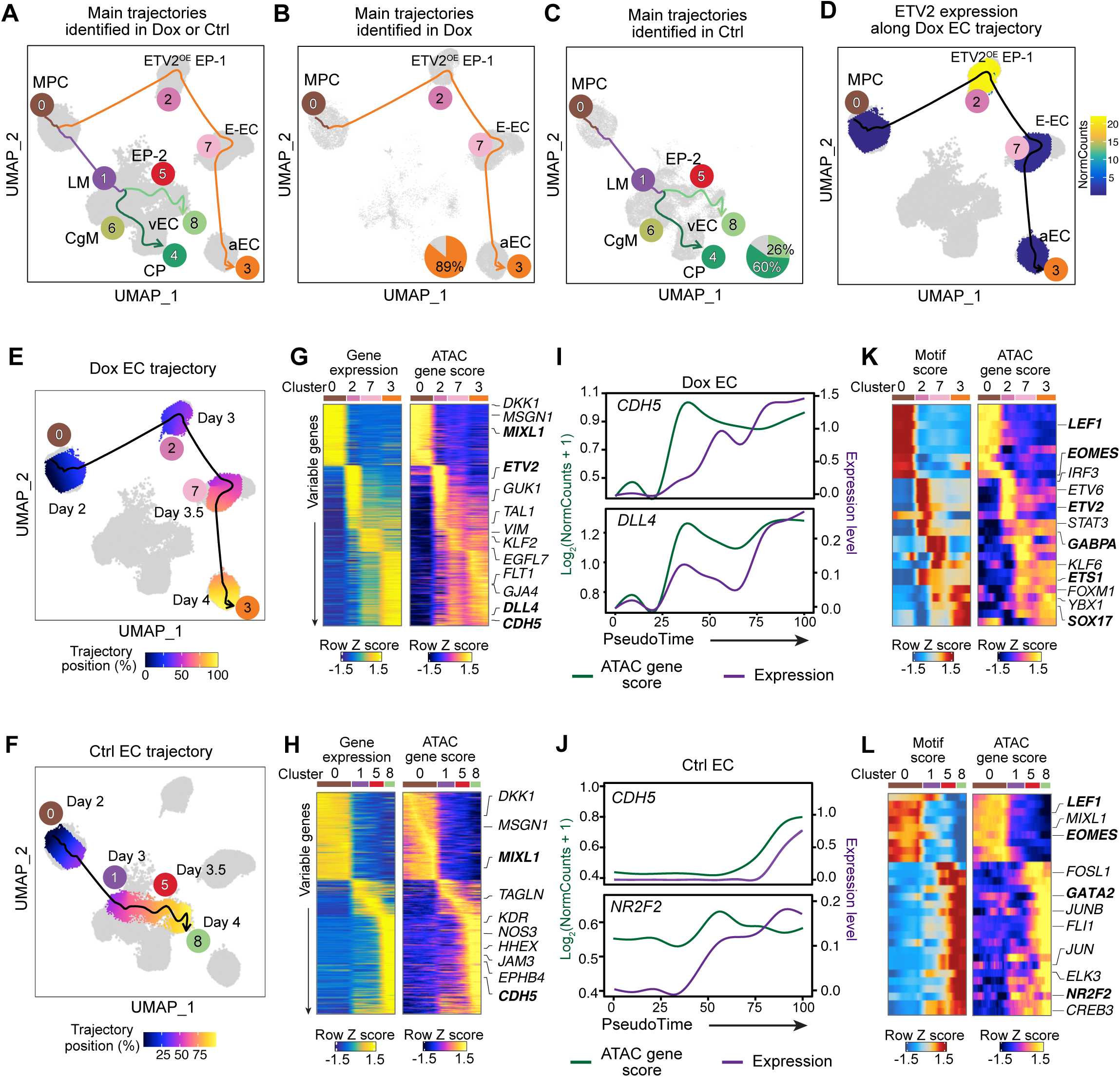
Reconstruction of differentiation trajectories with and without ETV2 overexpression. (**A-C**) UMAP of scATAC-seq cells highlighting the dominant trajectories identified in Dox (B) and Ctrl (C) groups. The cell clusters correspond to those in Fig. 2F. Pie chart in B,C showing the percentage of cells on the trajectory in indicated cluster at day 4. **(D)** Single-cell *ETV2* expression across pseudotime in the Dox EC trajectory. **(E,F)** UMAP visualization of the trajectory of MPC differentiation with (E) or without (F) ETV2 overexpression. Each cell is colored by the pseudotime and each cluster is annotated by sample collection time. The smoothed arrow represents a visualization of the interpreted trajectory in the UMAP embedding. **(G,H)** scRNA-seq gene expression (left) and scATAC-seq gene score (right) along the differentiation trajectory as a function of pseudotime. Selected genes had the greatest variation along the trajectory. Cell clusters are labeled according to their position along the pseudotime. **(I,J)** Gene expression and chromatin-derived gene accessibility score (ATAC gene score) dynamics of *CDH5* and *DLL4* in the Dox EC trajectory (I) or *CDH5* and *NR2F2* gene in Ctrl EC trajectory (J) across pseudotime. **(K,L)** Heatmap of TF regulators for which ATAC gene score is positively correlated with chromVAR TF deviation (motif score) across the Dox EC trajectory (K) or Ctrl EC trajectory (L). Cell clusters are labeled according to their position along the pseudotime.

We calculated the pseudotime position of each cell along the Dox or Ctrl EC trajectories (Fig. 3E,F) and used this to analyze the dynamics of gene expression and chromatin accessibility during the differentiation process (Fig. 3G,H). In both trajectories, the expression of the mesoderm marker *MIXL1* decreased with pseudotime, as did its ATAC gene score based on linked ATAC peaks. In the Dox trajectory, *ETV2* peaked at D3, followed by subsequent upregulation of pan-EC gene *CDH5* and then aEC gene *DLL4*^23^ (Fig. 3G, I and Extended Data Fig. 4A). Compared to the Dox trajectory, *ETV2* was only weakly upregulated at D3.5 in the Ctrl trajectory that terminated on vECs (Fig. 3F and Extended Data Fig. 4A), and *CDH5* upregulation occurred more weakly and later in pseudotime (Fig. 3H, J). The Ctrl trajectory was also characterized by upregulation of *NR2F2*, a TF important for vEC differentiation^24^ (Fig. 3J). A parallel analysis of the Ctrl trajectory that terminated on cardiac progenitors demonstrated upregulation of cardiac progenitor genes at late pseudotimes, and no induction of *ETV2* and *CDH5* (Extended Data Fig. 4B-E). Notably, across these analyses chromatin accessibility changes generally preceded gene expression changes in pseudotime, suggesting that activation of cis-regulatory elements was followed by later gene upregulation.

To gain insights into TFs driving gene expression and chromatin accessibility changes along Dox and Ctrl trajectories, we identified TFs with strongly correlated ATAC gene score and chromatin-based motif activity (motif score) across the Dox (Fig. 3K) or Ctrl (Fig. 3L and Extended Data Fig. 4E) trajectories.

TFs with high ATAC gene score and motif activity in MPCs (CL0) and lower values at later pseudotimes included mesoderm markers *LEF1* and eomesodermin (*EOMES)* (Fig. 3K,L). In the Dox trajectory, ETV2 motif activity and ATAC gene score sharply increased in ETV2^OE^ EP-1 and declined subsequently (Fig. 3K). The motif score of several other ETS-family TFs including *GABPA* and *ETS1* showed similar patterns. The motif accessibility and ATAC gene score of *SOX17*, an aEC-associated TF^25^, rapidly increased in cells belonging to the aEC cluster (CL3). In the Ctrl EC trajectory, the signature of activation of *ETV2* and other ETS-family TFs at intermediate pseudotimes was not clearly evident. The angiogenic GATA-family TF *GATA2*^26,27^ demonstrated increased motif activity and ATAC gene score at later pseudotimes, along with a signature for increased activity of the vEC TF *NR2F2* (Fig. 3L). In the Ctrl cardiac progenitor trajectory, at late pseudotimes we observed increased motif accessibility and ATAC gene score for cardiac progenitors markers, such as *HAND2* and *MEF2C* (Extended Data Fig. 4D,E).

Together, these analyses show that under Ctrl conditions, most cells follow a trajectory to other mesoderm fates including cardiac progenitors. MPCs undergo relatively reduced differentiation into endothelial progenitors and finally to ECs with venous features. In contrast, forced ETV2 expression in MPCs suppresses alternative fates and drives highly efficient differentiation to endothelial progenitors, which subsequently differentiate primarily into ECs with arterial features.

### TF regulatory networks that orchestrate EC differentiation

To identify the cell-type-resolved TFs that orchestrate EC differentiation, we analyzed the scATAC-seq data to identify differentially accessible peaks and the TF motifs enriched among differential peaks in each cell cluster (Fig. 4A). We further used scRNA-seq data to perform single-cell gene regulatory network analysis using SCENIC^28^. This identified the top regulons for each cluster, where each regulon is the gene network regulated by a TF (Fig. 4B).

**Fig. 4.**
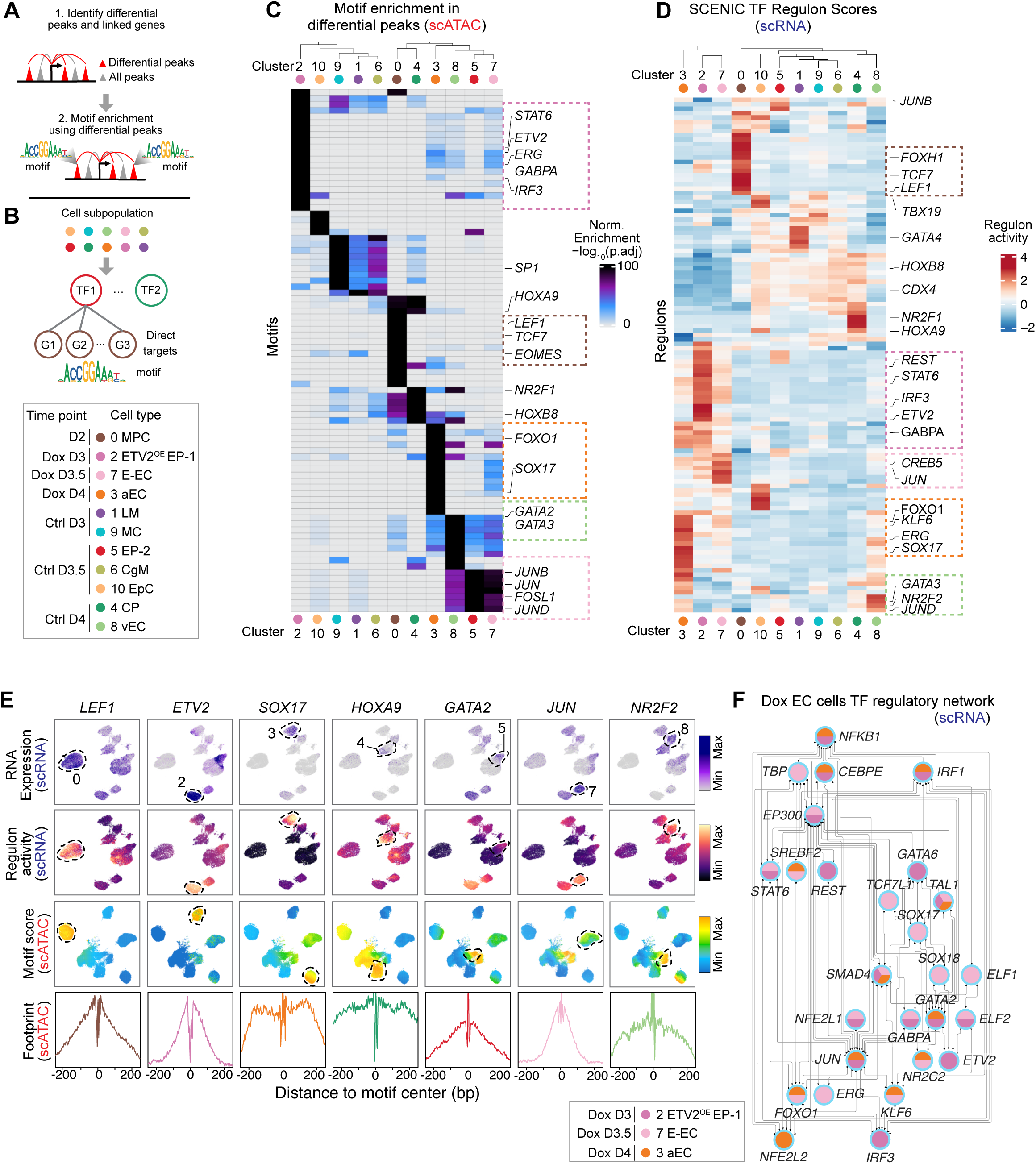
Integrative scATAC-seq and scRNA-seq analyses nominate cell-type-resolved TFs that regulate EC specification. (**A**) Schematic for identifying differential peaks and performing motif enrichment within differential peaks using scATAC-seq datasets. **(B)** Schematic for identifying TF regulons within different cell clusters using scRNA-seq datasets. **(C)** TF motif enrichment in differentially accessible peaks across each k-means cluster. Color indicates the motif enrichment based on the hypergeometric test. Selected TFs are labeled within dotted boxes that are colored to match the corresponding cell cluster. **(D)** Heatmap of SCENIC TF regulon activity across all cell clusters, showing cell-type specific transcriptional regulatory patterns. Selected TFs are highlighted within dotted boxes as in panel C. **(E)** Representative TFs with cell type selective regulatory activity. Row 1: TF expression plotted on scRNAseq UMAP; Row 2: regulon activity plotted on scRNAseq UMAP; Row 3: chromVAR TF deviation (motif score) plotted on scATACseq UMAP; Row 4, Tn5 bias-adjusted TF footprints. Gene expression was positively correlated with motif score across differentiation, representing positive TF regulators of differentiation. Lines in TF footprints are colored by the clusters shown in C,D. **(F)** Regulatory network of TFs in Dox clusters. Nodes represent TFs enriched in these clusters by motif enrichment (C) or SCENIC analyses (D). Edges represent protein-protein associations from STRING. Node colors indicate Dox EC clusters.

Motifs enriched in a cluster’s differentially accessible peaks identified candidate TFs that regulate gene expression and chromatin accessibility in the cluster. For example, the binding motifs for *EOMES* and *LEF1*, well established to regulate mesoderm formation^29,30^, were overrepresented in MPCs (CL0; Fig. 4C). ETV2^OE^ EP-1 (CL2) had a distinct TF motif profile characterized by enrichment in ETS-family binding motifs, including *ETV2* and *GABPA*, as well as motifs for interferon regulatory transcription factor (IRF) family members such as *IRF3* and STAT family members (Fig. 4C). The EP-2 cluster in Ctrl cells (CL5) and the early EC cluster in Dox cells (CL7) shared motif enrichment patterns. These clusters lacked ETS-family motifs and instead were highly enriched for AP-1 TFs such as *JUN* and *FOSL1*, and moderately enriched for GATA-family motifs. The motifs for *FOXO1* and *SOX17*, established aEC TFs, were enriched in aEC (CL3) (Fig. 4C).

Complementary to the motif analysis, analysis of the expression of TFs and their predicted downstream genes by SCENIC identified top TF regulons within each cluster (Fig. 4D). Top-ranked regulons included *ETV2*, *GABPA*, *REST*, and *STAT6* in ETV2^OE^ EP-1 (CL2); *JUN* and *CREB5* in early ECs (CL7) and *JUNB* in EP-2 (CL5); *FOXO1*, *KLF6*, and *SOX17* in aECs (CL3); and *NR2F2* in vECs (CL8) (Fig. 4D). By comparing the concordance between TF RNA expression level, TF regulon activity, and TF motif activity and footprint, we corroborated the regulatory activity of *LEF1* in MPCs (CL0), *ETV2* in ETV2^OE^ EP-1 (CL2), *JUN* in early ECs (CL7), *GATA2* in EP-2 (CL5), *SOX17* in aECs (CL3), and *NR2F2* in vECs (CL8) (Fig. 4E and Extended Data Fig. 5A).

We used the top-ranked TFs from motif-based and regulon analyses in Dox EC clusters (CL2, CL7, and CL3 in Fig. 4C,D) and protein-protein data from STRING^31^ to construct the TF-target network that regulates EC differentiation (Fig. 4F). Two prominent TFs in ETV2^OE^ EP-1 were *ETV2* and *GABPA*. These TFs shared several targets, including genes upregulated in Dox cells and associated with vessel structure and physiological function such as *RASIP1*, *RASGRP2,* and *ARHGEF2*) (Extended Data Fig. 5B).

Collectively, this analysis identified candidate TFs and TF-regulated targets in baseline and ETV2-induced MPC differentiation, including key TFs predicted to regulate EC specification.

### ETV2 modulates dynamic chromatin accessibility and enhancer remodeling

Given the divergence in chromatin accessibility and gene expression between basal and ETV2-directed MPC differentiation, and the function of ETV2 as a pioneer factor that opens chromatin to direct EC specification and differentiation, we analyzed chromatin accessibility changes in Ctrl vs. Dox conditions. To focus on the direct effects of ETV2, we focused on changes between D2 MPCs (CL0) vs. D3 control (lateral mesoderm or mesenchymal cell, CL1 and CL9) or D3 Dox (CL2, ETV2^OE^ EP-1; Fig. 5A,B). Most differentially accessible regions were accessible at both D2 and D3 and were referred to as “primed” accessible regions. Maintaining the accessibility of the large majority of these regions did not require forced ETV2 expression (Fig. 5A). In contrast, forced ETV2 expression opened 32,633 “de novo” regions in Dox condition that were inaccessible at D2, and the gain in accessibility of 97% of these regions required ETV2 overexpression. Forced ETV2 expression also closed 32,906 regions that were accessible at D2, and 64% of these closed regions were dependent on ETV2 overexpression.

**Fig. 5.**
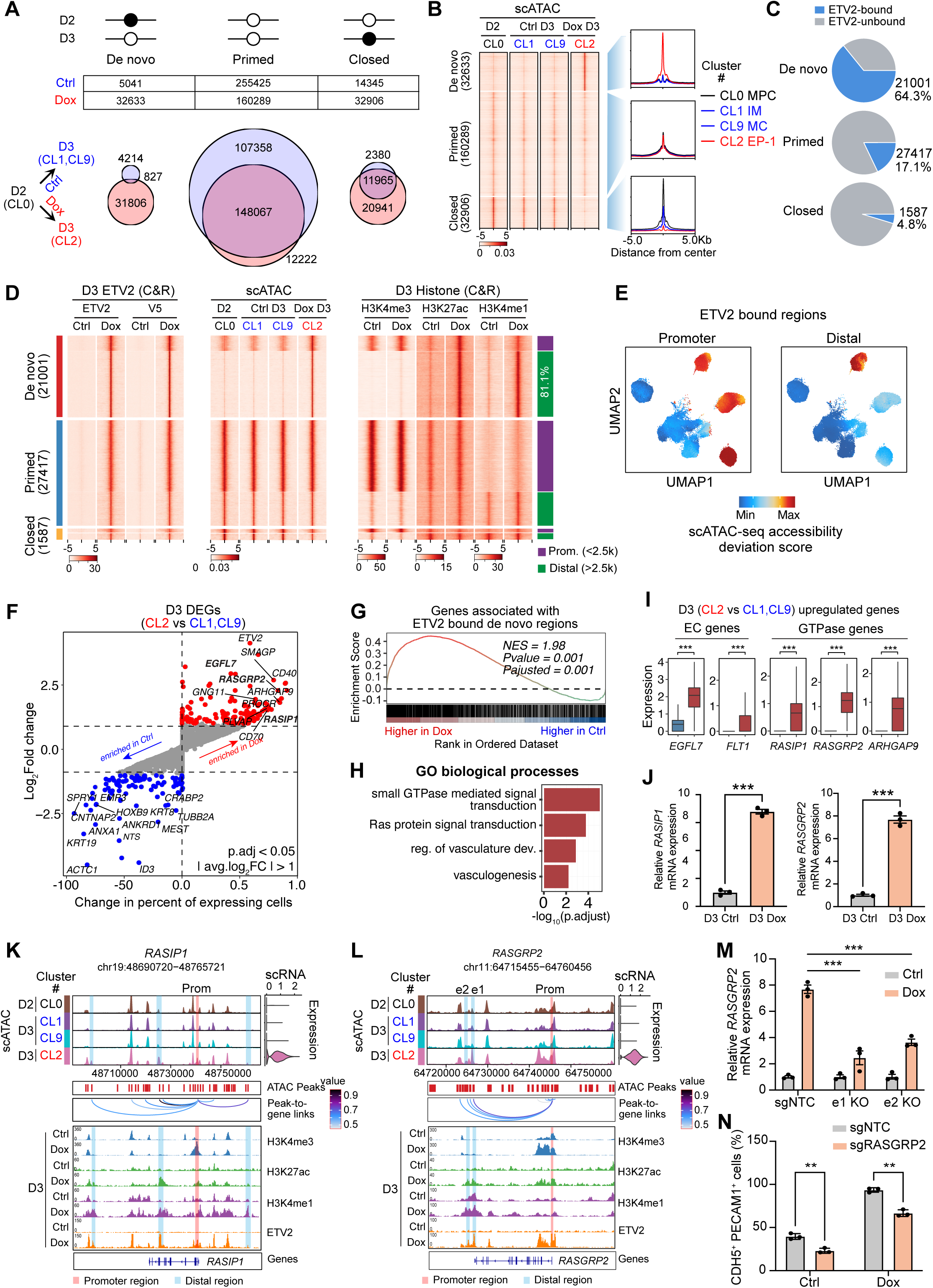
ETV2 promotes EC specification by modulating EC gene expression and GTPase signaling. (**A**) Regions that were differentially accessible between D2 (CL0 [MPC]) and D3 (Dox: CL2 [ETV2^OE^ EP-1]; Ctrl: CL1 [LM, Lateral mesoderm] and CL9 [MC, mesenchymal cell]) were classified into de novo (gained accessibility), primed (persistent accessibility), and closed (lost accessibility) regions. **(B)** Heatmap of the scATAC-seq signal for the indicated clusters and region classes (de novo, primed, closed). **(C)** Pie chart showing the percentage of ETV2 bound and unbound sites in each region classes (de novo, primed, closed). **(D)** Heatmap showing signal for the indicated chromatin features at all high confidence ETV2 binding sites. Regions are group by region class (de novo, primed, closed), and location with respect to TSSs (promoter, distal). **(E)** Accessibility of ETV2 bound promoter (<2.5 kb from TSS) and distal regions (>2.5 kb to TSS) during EC specification from MPCs. Accessibility deviation scores were mapped onto scATAC seq UMAP plots. **(F)** Genes differentially expressed between D3 Dox (CL2) and D3 Ctrl (CL1,9) cells. For each gene, the average log_2_fold change and difference in the percentage of cells with detectable expression are compared. Genes highlighted in red or blue have adjusted p values < 0.05 and | log_2_fold change | >1. **(G)** Gene-set enrichment analysis of genes linked to ETV2-bound de novo accessible regions. These genes were enriched among differentially expressed genes in D3 Dox compared to D3 Ctrl. **(H)** GO term analysis of D3 Dox upregulated genes compared to D3 Ctrl. **(I)** Expression of representative upregulated genes in D3 Ctrl and Dox scRNA data. **(J)** RT-qPCR analysis of *RASIP1* and *RASGRP2* mRNA expression in each group on D3. Two-tailed unpaired Student’s t-test. n=3. **(K,L)** Browser track showing chromatin features across clusters for the *RASIP1* (K) and *RASGRP2* (L) loci. Peak-to-gene loops are based on the correlation between peak accessibility and gene expression. Gene expression is shown for each cluster from scRNA-seq. **(M)** RT-qPCR measurement of *RASGRP2* expression after Cas9-mediated deletion of enhancer 1 (e1) and enhancer 2 (e2). Two-way ANOVA. n=3. **(N)** Quantification of the percentage of CDH5^+^ PECAM1^+^ cells generated at day 4 by differentiation of *RASGRP2* knockout or sgNTC iPSCs under Ctrl and Dox conditions. Two-tailed unpaired Student’s t-test. n=3. Data are mean ± s.e.m. **, P<0.01. ***, P<0.001. NTC, non-targeting control.

To investigate how ETV2 induces chromatin and enhancer remodeling to drive endothelial specification and differentiation, we assessed ETV2 binding sites in D3 Dox cells (Extended Data Fig. 6A). ETV2 chromatin occupancy was measured by CUT&RUN^32^ using ETV2 and V5 tag antibodies. We identified 43,441 and 95,276 genomic regions with enriched signals, respectively, with high reproducibility between biological replicates (Extended Data Fig. 6B). 43,284 high confidence peaks shared between both replicates and between ETV2 and V5 antibodies were defined as ETV2 regions (Extended Data Fig. 6C,D and Supplementary Table 4). ETV2 occupancy was barely detectable in D3 Ctrl cells (Extended Data Fig. 6D,E), consistent with little ETV2 expression at this time point and demonstrating the specificity of the chromatin occupancy signal. The majority of ETV2 regions were located in introns or distal intergenic regions (Extended Data Fig. 6F). Although promoters constitute a small fraction of the genome, they were strikingly enriched for ETV2 occupancy. Genome browser views of known ETV2 target genes *TJP2*, *ELK3*, *LMO2,* and *FLT1* demonstrated representative ETV2 binding at their enhancers and promoters (Extended Data Fig. 6G). In Dox cells, the majority (21,001, 64.3%) of de novo peaks were associated with ETV2 binding, demonstrating the pioneering activity of ETV2 to bind and open previously inaccessible chromatin sites. In comparison, much smaller fractions of primed (27,417, 17.1%) and closed (1,587, 4.8%) regions were ETV2-bound (Fig. 5C). The *ETV2* motif was the top-ranked motif among each of these ETV2 region subsets (Extended Data Fig. 6H). In addition to the ETV2 motif, we found a significant enrichment for binding motifs corresponding to ETS member *GABPA* and TFs from other families, including *STAT6*, *REST*, and *FOXM1* (Extended Data Fig. 6H). The *ETS-FOX* heterodimer motif^33^ was also highly enriched, particularly among de novo ETV2 regions. This motif was previously found to be highly enriched in p300-bound EC enhancers in vivo^34^ and to be sufficient to drive enhancer EC activity during embryonic vascular development^35^. *IRF3* enrichment was likewise strongly biased towards de novo compared to primed or closed ETV2 regions, suggesting that both *FOX* and *IRF3* can promote ETV2 pioneering activity. In contrast, *KLF6* were more strongly enriched among primed or closed compared to de novo regions, suggesting that these TFs do not participate in ETV2 pioneering activity.

We used CUT&RUN to assess ETV2 regions for H3K4me3, H3K27ac and H3K4me1, post-translational histone modifications associated with active promoters or enhancers^36,37^. Among de novo accessible ETV2 regions, ETV2 pioneer binding increased these marks on neighboring nucleosomes (Fig. 5D), suggesting the formation of active enhancers or promoters. Most de novo ETV2 regions (81.1%) were distributed at distal intergenic regions (Fig. 5D). At the same time, de novo ETV2 regions were highly enriched in promoters, given that promoters constitute a small fraction of the genome. Distal ETV2 regions were transiently accessible in ETV2^OE^ EP-1, whereas promoter ETV2 regions retained accessibility even Dox withdrawal (Fig. 5E). This suggests that ETV2-driven establishment of enhancers instructs early steps in EC specification, whereas ETV2-associated promoter activation is more stable through EC differentiation.

Genes neighboring de novo accessible ETV2 regions were enriched for functional terms related to regulation of small GTPase activity and regulation of blood vessel endothelial cell migration (Extended Data Fig. 6I). Unlike de novo accessible regions, closed regions were depleted for ETV2 occupancy (fraction of ATAC regions bound by ETV2: 4.8% closed, Fig. 5C). Genes associated with closed regions were generally downregulated between D2 and D3 (Extended Data Fig. 6J,K). TF motifs enriched in closed regions did not belong to the ETS family, and instead included TFs such as *LEF1* and *TCF* that are implicated in mesoderm specification^29,38^(Extended Data Fig. 6 L). These data suggest that ETV2-induced inactivation of many of these enhancers and genes was indirect.

Taken together, these results demonstrate ETV2’s pioneering activity to reshape the open chromatin landscape and identify enhancers and promoters directly regulated by ETV2.

### ETV2-regulated enhancers and target genes

To investigate ETV2 regulation of gene expression, we compared gene expression between D3 control (CL1 and CL9) and D3 Dox (CL2; Fig. 5F and Supplementary Table 5). Genes that neighbored de novo ETV2 regions were generally up-regulated (Fig. 5G). Differentially expressed genes (DEGs) that were up-regulated in the Dox group were highly enriched for genes related to GTPase activity and vasculature development (Fig. 5H,I). For example, ETV2 overexpression activated *EGFL7* and *FLT1*, which are required for angiogenesis, EC migration, and endothelial integrity^39,40^. In contrast, down-regulated DEGs were enriched for functional terms related to embryonic organ development, cardiac ventricle development and muscle cell differentiation (Extended Data Fig. 7). These results suggest that ETV2 acts as a transcriptional activator to promote EC differentiation, vascular structure, and EC function.

*RASIP1* (Ras interacting protein 1), a Rho GTPase activity regulator that controls cell adhesion and junctional stabilization^41^. *RASGRP2* (RAS guanyl-releasing protein 2) acts as a guanine nucleotide exchange factor and activates small GTPase RAP1^42^. We confirmed the upregulation of GTPase regulators *RASIP1* and *RASGRP2* in Dox cells by RT-qPCR (Fig. 5J). We also identified candidate cis-regulatory elements (CREs) that might regulate these genes, based on ETV2 binding, ETV2-dependent enhancer features (chromatin accessibility, H3K27ac, and H3K4me1), and strong peak-to-gene links predicted based on correlation between region accessibility and gene expression (Fig. 5K,L). To test the requirement for the predicted CREs to activate these genes, we selected two adjacent CREs to *RASGRP2* (e1 and e2; Fig. 5L) for Cas9-mediated deletion. Deletion of either e1 or e2 strongly attenuated *RASGRP2* upregulation by Dox at D3 (Fig. 5M). Furthermore, we confirmed that Cas9-mediated *RASGRP2* knockout (sgRASGRP2) reduced EC specification efficiency in both Ctrl and Dox conditions (Fig. 5N).

Taken together, these results show that ETV2 promotes EC specification by activating target genes through binding to distal regulatory elements. We identified genes and candidate CREs regulated directly by ETV2 (Supplementary Table 3). We validated two ETV2-dependent enhancers of *RASGRP2*, and showed that this gene is required for efficient EC specification.

### CRISPR screening uncovers regulators of EC specification

The analyses of regulons and TF motifs involved in EC specification and differentiation (Fig. 4) and of direct ETV2 target genes (Fig. 5) nominated transcriptional regulators that may collaborate with ETV2 to drive EC lineage commitment. To experimentally assess the contribution of these 511 candidate genes (Supplementary Table 6) to EC lineage commitment induced by ETV2, we performed a focused, pooled CRISPR screen^43^. We synthesized an sgRNA library containing 6 independent guides per gene, along with 90 negative control sgRNAs that do not target the human genome, and cloned them as a pool into a lentiviral vector that expresses mCherry (Supplementary Table 6). We transduced TRE3G-ETV2 iPSCs with Lenti-Cas9-GFP and the Lenti-sgRNA-mCherry library lentivirus (Fig. 6A). GFP and mCherry double-positive cells were sorted and differentiated under Dox conditions. At D4, CDH5^+^ PECAM1^+^ cells were isolated by FACS. Next-generation sequencing of single guide RNAs (sgRNAs) within the double-positive ECs were compared with input to identify genes targeted by sgRNAs that were significantly enriched or depleted (Fig. 6A).

**Fig. 6.**
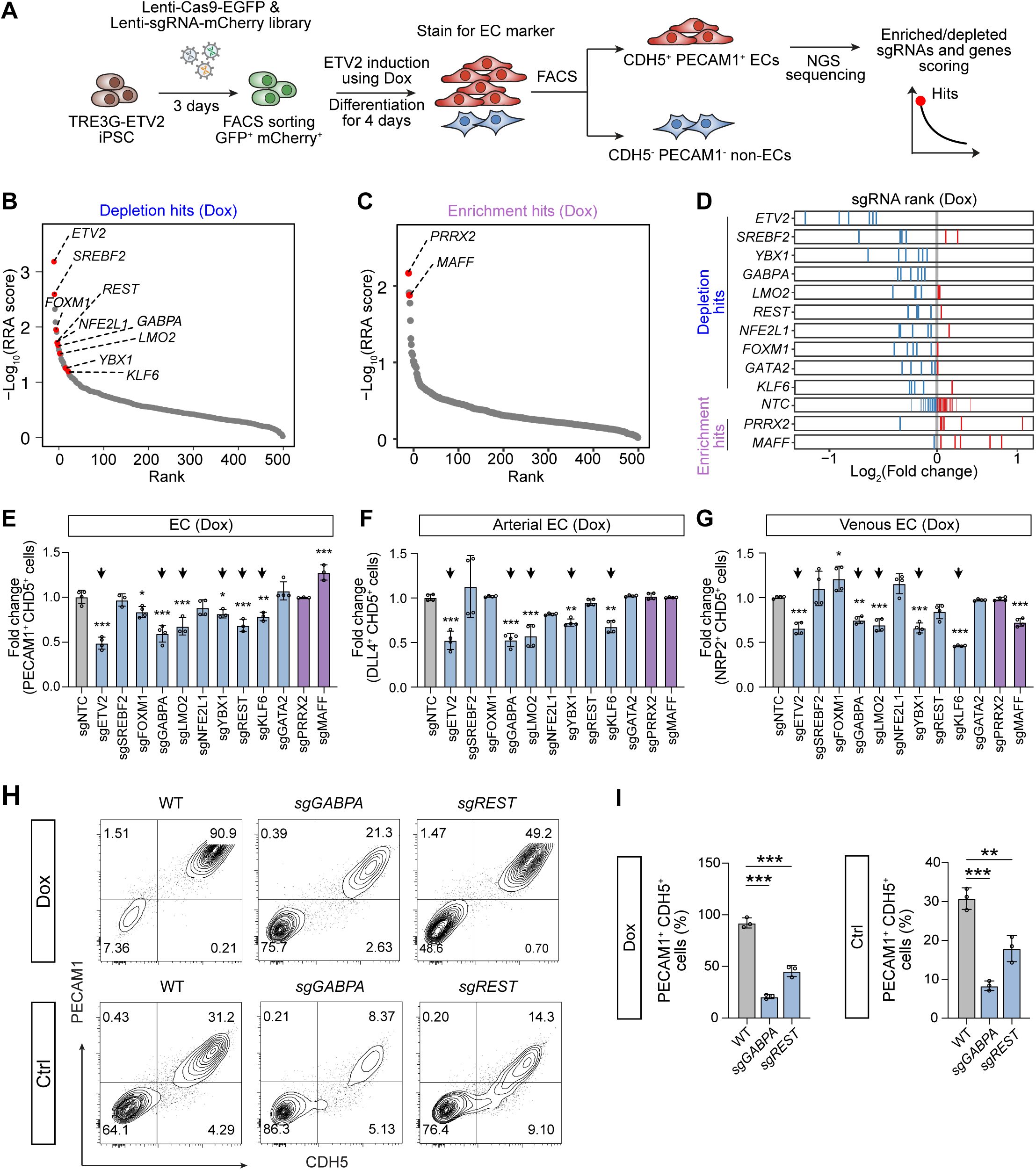
CRISPR-Cas9 screening identifies GABPA and REST as key regulators of ETV2-drived EC specification. (**A**) Schematic of the CRISPR screening approach for identifing TF regulators of EC specification. Oligonucleotide synthesis was used to generate a pooled Lenti-sgRNA-mCherry library containing 3156 gRNAs that targeted candidate transcriptional regulators. TRE3G-ETV2 iPSCs were transduced with lenti-Cas9-GFP and the lenti-sgRNA-mCherry library to establish stably transduced cell pools. The cells were then differentiated with or without Dox. FACS for CDH5 and PECAM1 was used to separate ECs from non-ECs. Enriched and depleted sgRNA were identified by next-generation sequencing (NGS) of sgRNAs. **(B,C)** Depletion hits (B) and enrichment hits (C). Each gene was ranked based on the MAGeCK robust ranking aggregation (RRA) score. The y axis represents – Log_10_(RRA score). The selected top hits are labeled in red. **(D)** Top screening candidate genes, showing relative rank for each individual sgRNA (6 sgRNAs/gene). The x axis represents log_2_(fold change). Blue and red bars indicates sgRNA depletion or enrichement, respectively. **(E-G)** Validation of screening hits. iPSCs were transfected with a non-targeting control sgRNA (sgNTC) or sgRNAs targeted to the indicated gene. Differentiation efficiency under Dox conditions to PECAM1^+^ CDH5^+^ ECs (E), DLL4^+^ CDH5^+^ aECs (F), or NRP2^+^ CDH5^+^ vECs (G) was measured by flow cytometry. n = 3 to 4. **(H,I)** Flow cytometry analysis (H) and quantification (I) for percentage of PECAM1^+^ CDH5^+^ cells in WT, *GABPA* knockout and *REST* knockout group under Ctrl and Dox condition on day 4. One-way ANOVA with Dunnett’s multiple comparison test to WT control. n = 3. Data are mean ± s.e.m. *, P<0.05; **, P<0.01; ***, P<0.001. ns, not significant. NTC, non-targeting control.

Overall, the screen identified genes whose knockout significantly inhibits (depleted hits) or promotes (enriched hits) EC differentiation induced by ETV2 (Fig. 6B,C). We selected 10 depleted and 2 enriched hits for individual validation. For each gene, we transduced TRE3G-ETV2 iPSCs with Lenti-Cas9-GFP and a Lenti-sgRNA-mCherry that expressed the gene’s 3 most significant sgRNAs (Fig. 6D). We did not observe changes in pluripotency marker SSEA-4 in the CRISPR-treated iPSCs compared to non-targeting control (sgNTC) (Extended Data Fig. 8A). We next tested the CRISPR-treated iPSCs pools for EC differentiation under Dox. Six of the 12 tested genes significantly altered EC specification efficiency (Fig. 6E). Knockout of *LMO2, KLF6, YBX1, GABPA*, and *REST* impaired EC specification efficiency, whereas knockout of *MAFF* increased it. These five genes required for efficient EC specification were also required for efficient differentiation of both aECs and vECs (Fig. 6F,G), suggesting a potential role in EC specification.

We also investigated the regulation of EC specification and differentiation in Ctrl conditions. Under these conditions, we likewise observed that *KLF6, YBX1, GABPA*, and *REST* were required for efficient EC differentiation and that *MAFF* limited EC differentiation efficiency (Extended Data Fig. 8B-D). These results suggest that these genes also participate in EC specification at endogenous ETV2 expression levels.

Among these genes, *LMO2*, a known ETV2 downstream target^8^, is an established regulator of EC development^44^. *KLF6* regulates EC remodeling and response to injury, and *KLF6* knockout mice show defects in yolk sac vascularization^45^. *YBX1*, a DNA and RNA binding factor, has been implicated in tumor angiogenesis^46^. *GABPA*, an ETS-family transcription factor that is required for early embryogenesis, has not been functionally analyzed in EC specification or differentiation. Similarly, the function of *REST*, a well-characterized transcriptional repressor^47,48^, in EC specification or differentiation has not been explored. Then, we generated knockout cells of *GABPA* and *REST* with high editing efficiency in the ETV2-inducible iPSCs to further characterize their roles in EC differentiation. We differentiated these iPSCs to MPCs and did not observe a significant change in mesoderm marker genes (Extended Data Fig. 8E), indicating that mesoderm formation was not affected. Consistent with the results from CRISPR-treated cell pools, *GABPA* and *REST* knockout cells yielded a significantly reduced fraction of PECAM1^+^ CDH5^+^ ECs on D4 compared with the parental cells in either Dox or Ctrl conditions (Fig. 6H,I).

Examination of the expression profile of these genes along the Dox EC trajectory (Fig. 3B) showed that *REST* and *GABPA* maintain expression levels during the peak of ETV2 expression (Extended Data Fig. 8F). TF motif discovery at de novo ETV2 regions recovered the TFs *GABPA* and *REST* (Extended Data Fig. 6H). This suggests that *GABPA* and *REST* may functionally interact with pioneer factor *ETV2* to open chromatin and promote EC lineage commitment.

Taken together, our loss of function screen and validation experiments identified GABPA and REST as functional regulators that may act with pioneer factor ETV2 in EC specification.

### GABPA and REST co-occupy ETV2 regions in ETV2^OE^ endothelial progenitor cells

To test the hypothesis that GABPA and REST regulate EC lineage commitment by co-occupying ETV2 regions, we measured their genomic occupancy using CUT&RUN on D3 of Dox-induced EC differentiation of iPSCs (Supplementary Table 4). Motif enrichment analysis of the CUT&RUN data recovered GABPA and REST binding motifs among the top two most significant motifs (Fig. 7A), demonstrating the high quality of data. Notably, ETV2 was the other top enriched motif in the GABPA– or REST-bound regions (Fig. 7A). Overlap with ETV2 regions revealed that ETV2 co-occupied 51% (17,007/32,942) of GABPA regions and 75% (23,779/31,583) of REST regions (Fig. 7B). Conversely, GABPA and REST co-occupied 39% and 55% of ETV2 regions, respectively. Nearly all of these peaks were within de novo and primed accessible chromatin regions of D3 ETV2^OE^ endothelial progenitor cells (Fig. 5B and Fig. 7C). Indeed, a significant fraction of the regions co-occupied by ETV2 and GABPA or REST were de novo accessible regions in D3 ETV2^OE^ endothelial progenitor cells (GABPA, 27%; REST, 37%). However, few GABPA or REST only peaks were within de novo accessible chromatin regions, indicating that GABPA and REST engage the open chromatin and have little independent pioneering activity. The CUT&RUN occupancy signal for ETV2, GABPA, and REST was substantially stronger at co-occupied regions compared to regions occupied by each TF alone (Fig. 7D,E), suggesting collaborative binding.

**Fig. 7.**
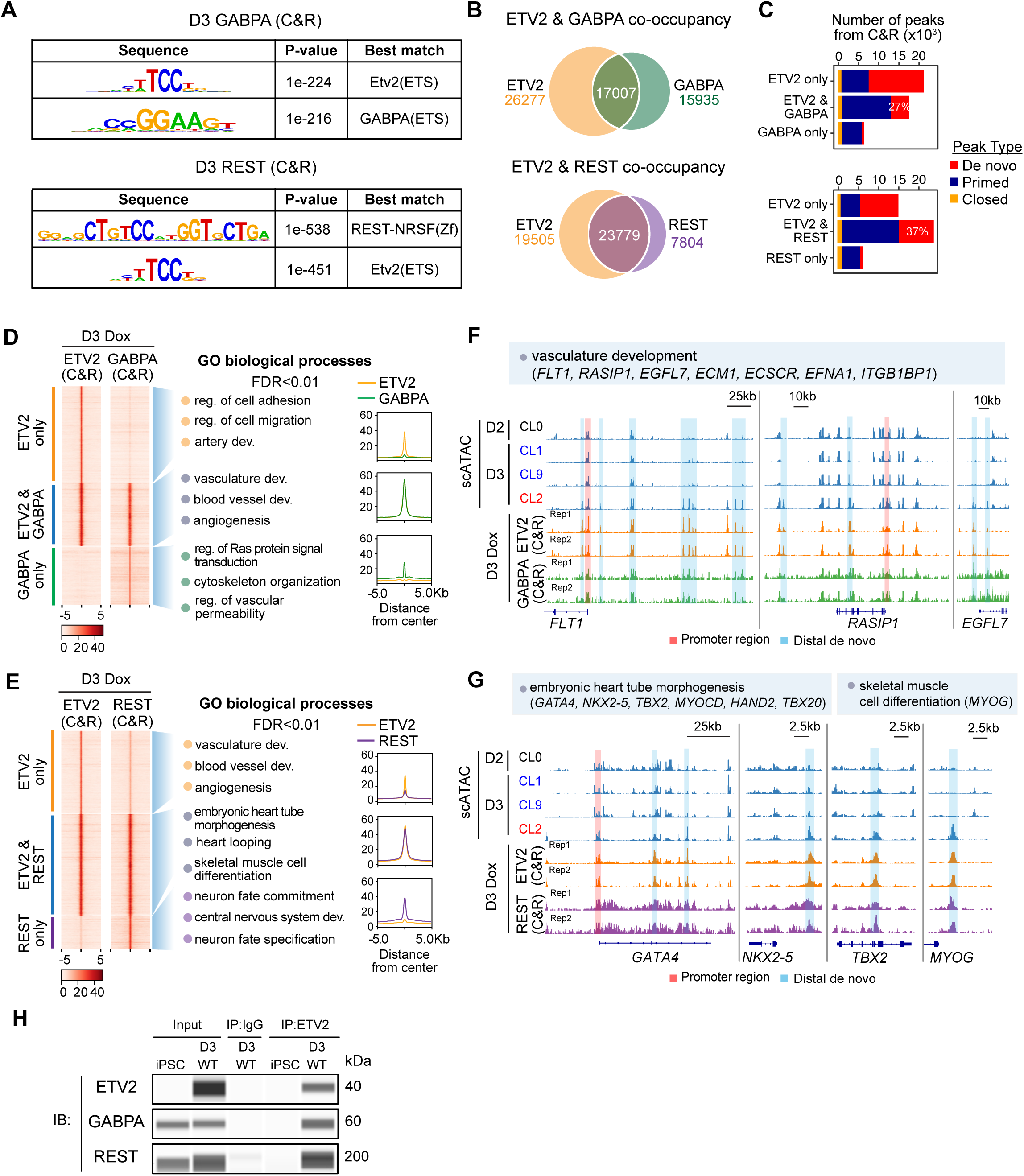
ETV2 recruits GABPA and REST to enhancers. (**A**) The two top enriched motifs in all GABPA regions or REST regions in D3 Dox cells. **(B)** Venn diagrams showing the overlap between GABPA and ETV2 binding peaks (top), or between REST (bottom) and ETV2 binding peaks in D3 Dox cells. **(C)** Binding of GABPA or REST within ETV2-bound peaks, separated into de novo, primed and closed subsets based on the scATAC-seq data presented in Fig. 5. **(D,E)** Left panel, heatmaps showing GABPA (D) or REST (E) binding signals at ETV2 regions, demonstrating ETV2 co-occupancy with GABPA or REST. Middle panel, regions with different TF occupancy patterns had distinct enriched GO biological processes terms (GREAT; FDR < 0.01). Right panel, mean CUT&RUN signals in each region. **(F,G)** Genome browser views showing scATAC-seq in D2 (CL0), D3 Ctrl (CL1,9), D3 Dox (CL2) cells, and GABPA (F), REST(G), ETV2 binding signals of representative regions in D3 Dox cells. De novo distal sites are shaded blue. **(H)** Co-immunoprecipitation (Co-IP) using ETV2 or IgG control antibodies, followed by capillary immunoblotting (IB) for ETV2, GABPA or REST, in iPSC or Dox D3 WT cells.

We analyzed the functional terms enriched among genes neighboring regions occupied by ETV2 and GABPA, ETV2 only, or GABPA only (Fig. 7D). ETV2-only regions were associated with the regulation of cell adhesion, cell migration, and artery development, GABPA-only regions were associated with cytoskeleton organization and regulation of vascular permeability, and co-bound regions revealed enrichment for ontologies involved in vasculature development, and angiogenesis. For example, GABPA and ETV2 co-occupied regions near several upregulated genes involved in vasculature development, such as *FLT1*, *RASIP1* and *EGFL7* (Fig. 7F), indicating that ETV2 and GABPA cooperate to participate in early vascular development. We found that GABPA and ETV2 co-occupied their promoters and enhancers, which were also opened following ETV2 overexpression in scATAC-seq data (CL2 vs CL1,9; Fig. 7F).

Interestingly, ETV2 and REST regions similarly displayed distinct functional categories between ETV2 plus REST, ETV2-only, and REST-only subsets (Fig. 7E). ETV2-only regions were enriched for GO terms related to vascular development, REST-only regions were related to neuron fate commitment and central nervous system development, and ETV2 plus REST co-occupied regions were associated with the embryonic heart tube morphogenesis, heart looping, and skeletal muscle cell differentiation. Among the REST and ETV2 co-occupied regions were de novo accessible regions of several cardiac (e.g., *GATA4*, *NKX2-5,* and *TBX2*) and muscle genes (e.g., *MYOG*; Fig. 7G), which were silenced in mature ECs, indicating that ETV2 recruits REST to suppress non-EC lineage genes.

We used co-immunoprecipitation assays to test whether GABPA or REST form a complex with ETV2 to regulate EC lineage commitment. We found that GABPA and REST both co-immunoprecipitated with ETV2 (Fig. 7H). These results demonstrate that ETV2 opens chromatin and recruits GABPA and REST to regulate target genes in ETV2^OE^ endothelial progenitor cells.

### GABPA and ETV2 collaboratively activate early EC genes

To unravel the contribution of GABPA to transcriptional regulation by ETV2, we performed bulk RNA-seq on WT TRE3G-ETV2 cells under Ctrl and Dox conditions at D2 and D3, and on D3 Dox *GABPA* knockout TRE3G-ETV2 cells (24 hours of ETV2 overexpression; Extended Data Fig. 9A). Strikingly, most genes upregulated in WT by ETV2 overexpression (D3 Dox WT vs. D2) were strongly downregulated in the *GABPA* knockout following ETV2 overexpression (intersection was 73% of D3 Dox WT upregulated genes and 67% of *GABPA* knockout downregulated genes; Fig. 8A, Extended Data Fig. 9B,C, and Supplementary Table 5). ETV2 and GABPA co-occupied most (76%) of these overlapping genes, supporting the requirement of GABPA for ETV2-mediated gene activation. *GABPA* knockout downregulated genes were enriched for GO terms related to the regulation of angiogenesis (*FLT1, HHEX*, and *ECM1*) and small GTPase-mediated signal transduction (*RASIP1*, *SIPA1,* and *RASGRP2*; Fig. 8B,C).

**Fig. 8.**
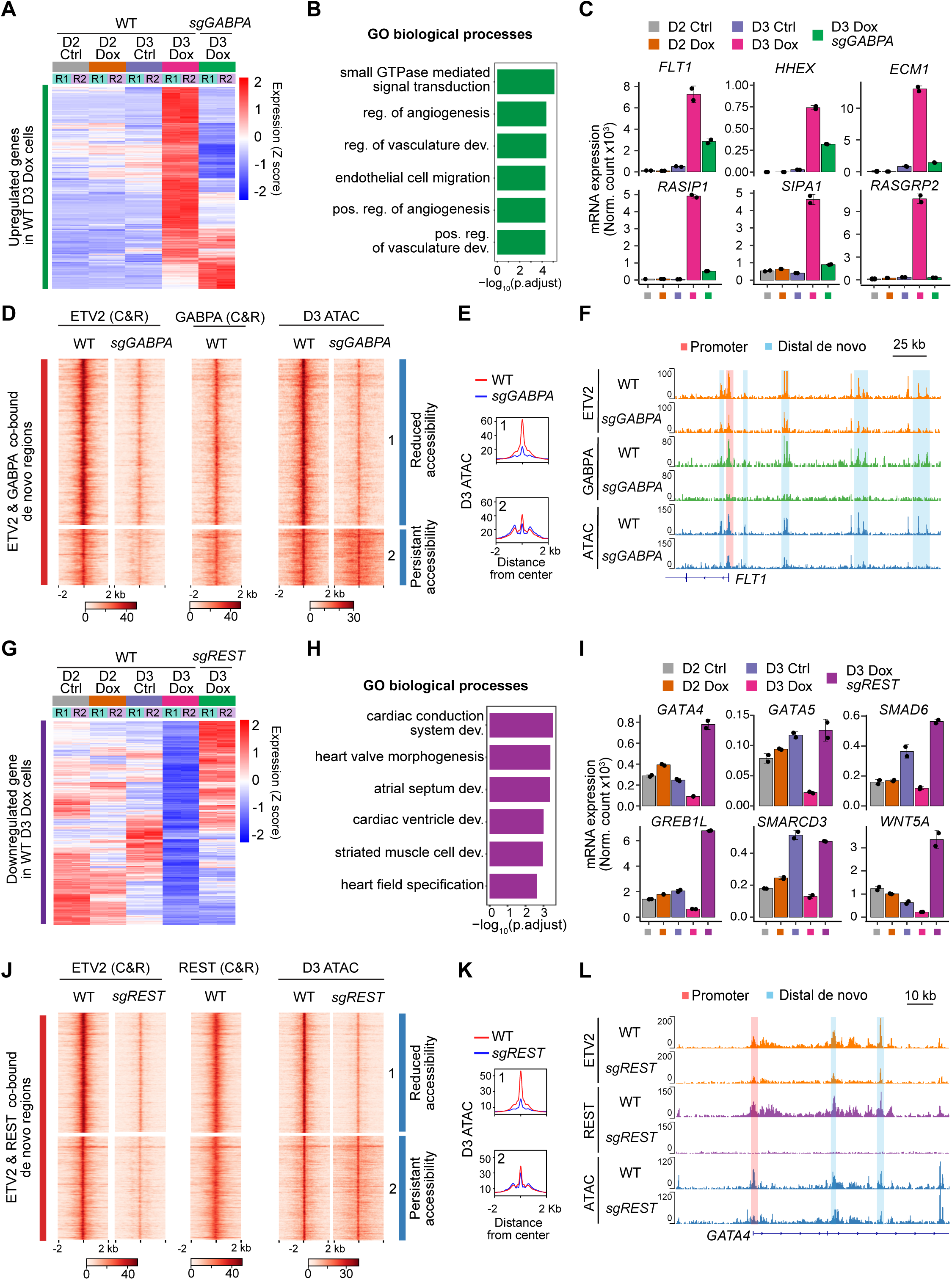
GABPA and REST promote EC specification by collaborating with ETV2 to open chromatin and regulate gene expression. (**A**) Expression of WT D3 Dox upregulated genes in each WT and *GABPA* knockout group. Each row is z-score normalized. R1 and R2 indicate biological replicates. **(B)** GO term analysis of genes that were downregulated in Dox D3 *GABPA* knockout cells. **(C)** Examples of genes downregulated in Dox D3 *GABPA* knockout cells by RNA-seq. n = 2. **(D,E)** Heatmaps (D) show indicated chromatin features from Dox D3 WT and *GABPA* knockout cells at ETV2 and GABPA co-bound de novo accessible regions (presented in Fig. 5D). Region subsets labeled 1 had reduced accessibility in *GABPA* knockout, and those labeled 2 had similar accessiliby in WT and *GABPA* knockout. Aggregation plots (E) show average D3 ATAC signal from region subsets in Dox D3 WT or *GABPA* knockout. **(F)** Representative genome browser view at *FLT1* showing ETV2 binding, GABPA binding and ATAC-seq in WT and *GABPA* knockout cells. **(G)** Expression of WT D3 Dox downregulated genes in each WT and *REST* knockout group. Each row is z-score normalized. R1 and R2 are biological replicates. **(H)** GO term analysis of genes that were upregulated in Dox D3 *REST* knockout cells. **(I)** Examples of genes upregulated in Dox D3 *REST* knockout cells by RNA-seq. n = 2. **(J,K)** Heatmaps (J) show indicated chromatin features for Dox D3 WT and *REST* knockout cells at ETV2 and REST co-bound de novo regions (presented in Fig. 5D). Region subsets labeled 1 had reduced accessibility in *REST* knockout, and those labeled 2 had similar accessiliby in WT and *REST* knockout. Aggregation plots (K) show average D3 ATAC signal from region subsets in Dox D3 WT or *REST* knockout. **(L)** Representative genome browser view at *GATA4* showing ETV2 binding, REST binding, and ATAC-seq in WT and *REST* knockout cells.

To investigate the effect of *GABPA* knockout on ETV2 occupancy and on chromatin accessibility, we conducted ETV2 CUT&RUN and ATAC-seq in WT and *GABPA* knockout cells at D3 Dox. The depletion of GABPA strongly reduced ETV2 binding in de novo peaks at D3 (Fig 5D, Fig. 8D and Extended Data Fig. 9D). This reduction occurred at regions with GABPA co-occupancy (Fig. 8D), indicating that GABPA regulates ETV2 pioneering activity directly. Consistent with altered ETV2 pioneering activity, GABPA depletion reduced accessibility of 46.5% of WT de novo ETV2 regions (Extended Data Fig. 9E). Among ETV2 and GABPA co-occupied de novo accessible regions, 75% had reduced accessibility upon depletion of GABPA (group 1, Fig. 8D,E). For example, we observed that in the depletion of *GABPA*, ETV2 occupancy decreased in many enhancers of *FLT1*, an angiogenic gene, and their chromatin accessibility decreased (Fig. 8F).

These results demonstrate that GABPA collaborates directly with ETV2 to shape the chromatin accessibility landscape of ETV2^OE^ endothelial progenitor cells and activate downstream endothelial genes.

### ETV2-dependent REST recruitment is required to repress non-EC genes

REST binds to a 21 bp consensus sequence known as repressor element 1 (RE1) and represses gene transcription^49^. To define which ETV2 target genes and their CREs are co-regulated by ETV2 and REST, we compared gene expression profiles of D3 Dox *REST* knockout TRE3G-ETV2 cells to WT TRE3G-ETV2 cells at D2 or D3 under Dox or Ctrl conditions (Extended Data Fig. 9F). Remarkably, many of the genes downregulated in WT during Dox-induced EC differentiation (D3 Dox WT vs. D2) were significantly upregulated in *REST* knockout cells (intersection was 72% of D3 Dox WT downregulated genes and 44% of *REST* knockout upregulated genes; Fig. 8G, Extended Data Fig. 9G,H, and Supplementary Table 5). ETV2 and REST co-occupied the majority (66%) of these overlapping genes. GO terms enriched among these genes were related to heart specification, and affected genes included *GATA4*, *GATA5*, *SMAD6*, *GREB1L*, *SMARCD3*, and *WNT5A*, all known to regulate heart development (Fig. 8H,I). These results show that ETV2 opens chromatin and recruits REST to repress the expression of non-EC lineage genes, especially cardiac genes, during EC differentiation. Cardiac differentiation was also a major alternative trajectory in Ctrl conditions (Fig. 3C), consistent with the need to actively repress this fate to achieve efficient EC specification.

We investigated the effect of *REST* knockout on ETV2 occupancy and chromatin accessibility by performing ETV2 CUT&RUN and ATAC-seq in D3 Dox WT and *REST* knockout cells. Further supporting the cooperation of REST and ETV2 during EC specification, *REST* knockout reduced ETV2 occupancy of de novo accessible regions at regions co-occupied by REST (Fig. 8J and Extended Data Fig. 9I). Reflective of reduced ETV2 pioneering activity, REST inactivation reduced the accessibility of 34.6% of de novo accessible ETV2 regions (Extended Data Fig. 9J). Among ETV2 and REST co-occupied de novo accessible regions, 53% had reduced accessibility in *REST* knockout (group 1, Fig. 8J,K). We examined the CREs of GATA4, a key regulator of cardiac differentiation. The ETV2 and REST co-occupied de novo GATA4 CREs were less accessible in the absence of REST and were no longer bound by ETV2 or REST (Fig. 8L). Loss of REST repression is consistent with GATA4 upregulation in *REST* knockout D3 Dox (Fig. 8I).

Together, our findings support the model that ETV2 pioneering activity is influenced by REST, and that ETV2 recruitment of REST is required to repress the cardiac lineage during EC specification.

## Discussion

To gain insights into mechanisms of cell specification, we developed a highly efficient system in which overexpression of the pioneer factor ETV2 in MPCs reproducibly drives EC specification in two days. Using this system, we established a single-cell transcriptome and accessible chromatin atlas to capture the transcriptional and epigenetic dynamics of EC specification. Using this resource, we demonstrated that ETV2 overexpression overcomes two key bottlenecks in EC specification: (1) formation of EC progenitors, and (2) diversion of cells into non-EC fates. We defined ETV2-bound de novo accessible regions, which depend upon ETV2 binding and pioneering activity to open chromatin, establish active enhancer marks, and activate EC genes. From these datasets, we nominated candidate TFs likely to collaborate with ETV2 to promote efficient EC specification. Functional CRISPR screening of these candidates followed by validation studies revealed several TFs with previously unrecognized roles in EC specification. Among these were the transcriptional activator GABPA and the transcriptional repressor REST.

Pioneer factors promote lineage specification by binding recognition sites within nucleosomal DNA, making the DNA accessible so that typical TFs can bind, recruit transcriptional activators, and stimulate the expression of genes that promote lineage commitment^3^. This canonical pioneering role of ETV2 was exemplified by its interaction with GABPA. *GABPA* expression level did not significantly increase along the EC trajectory, but following ETV2 induction it co-occupied many ETV2-bound de novo regions. This, together with its physical interaction with ETV2 and the loss of ETV2 occupancy in *GABPA* knockout, suggest that GABPA collaborates with and enhances ETV2 pioneering activity. This result complements and reinforces prior studies that demonstrated that typical TFs modulate the binding affinity and specificity of pioneer TFs^50,51^. Although *ETV2* expression is transient during EC specification, GABPA may continue to occupy and activate ETV2-pioneered ETS motifs after *ETV2* down-regulation, since *GABPA* and *ETV2* both belong to the ETS TF family and share similar DNA binding motifs. ETV2 was previously shown to activate the expression of other angiogenic ETS family TFs such as *FLI1* and *ERG* in a feed-forward loop described as an “ETS switch”^8,27^. Here, ETV2 is not predicted to activate GABPA expression; rather, ETV2 and GABPA collaborate to enhance DNA binding and establish de novo chromatin accessibility.

Our study revealed two additional facets of ETV2 pioneering activity. First, we found that ETV2 promotes sustained chromatin accessibility of promoters, whereas its effect on accessibility of distal regions is transient. ETV2 regions, including de novo accessible regions, were over-represented for promoters, and ETV2 bound promoters tended to remain accessible in Dox cells through D4, well after ETV2 expression fell to basal levels. In contrast, accessibility of ETV2-bound distal regions tended to be transient, with many becoming inaccessible at D3.5 and D4. This suggests that pioneer factors like ETV2 may induce epigenetic memory by opening promoter regions, which then remain open independent of the pioneer factor. This persistent promoter accessibility may reflect the activation of lineage-specific transcriptional programs that sustain promoter activity in the absence of the initiating pioneer factor.

Second, we found that ETV2 recruits the transcriptional repressor REST to silence non-EC lineage genes, particularly cardiac genes. Originally discovered as a repressor of neuronal gene expression^52^, REST is widely expressed outside of the nervous system and contributes to stem cell pluripotency and differentiation^53,54^. In keeping with a prior scRNAseq study^55^, under control conditions without ETV2 overexpression, the majority of cells acquired cardiac and other non-EC fates, in part accounting for inefficient EC specification. Previous studies showed that ETV2 inactivation in mouse embryos and murine embryonic stem cells/embryonic bodies resulted in expansion of cardiac and other mesodermal lineages^56,57^. These data indicate that ETV2 balances mesodermal lineage commitment. Our study suggests that ETV2 both actively stimulates EC specification, through activators such as GABPA, and actively represses alternative lineage choices, through repressors such as REST. We showed that REST ablation reduced EC specification and upregulated cardiac genes. ETV2 did not activate REST expression; rather it physically interacted with REST and recruited it to ETV2-bound de novo accessible regions, where it repressed expression of alternative lineage genes. We propose that recruitment of transcriptional repressors may be a general feature of pioneer factors that limits transcriptional activation by some newly opened chromatin regions to suppress commitment to alternative lineages.

Early in vascular development, ECs adopt arterial or venous identities, each characterized by distinct transcriptional programs. Developmentally, aECs emerge slightly earlier than vECs in murine embryos^58^. Notch, high VEGF, and ERK signaling promote aEC differentiation, whereas PI3K and the transcription factor NR2F2 promote vEC differentiation^59,60^. We observed that high and early induction of ETV2 promoted a distinct EC specification pathway that terminated on ECs with arterial features, whereas lower and later endogenously expressed ETV2 yielded ECs with venous features. This observation suggests that the level or timing of ETV2 during EC specification may be an additional factor that biases arteriovenous differentiation. This possibility is consistent with high VEGF stimulating aEC differentiation, since VEGF induces ETV2 expression^61,62^. Additional experiments are required to further investigate the role of ETV2 in the regulation of the arteriovenous differentiation of EC progenitors.

Vascular regeneration is a promising therapeutic strategy for diseases that impair vascular function, including atherosclerosis and diabetes^63–65^. ETV2 overexpression reprograms iPSCs, iPSC-derived MPCs, and somatic cells such as fibroblasts into functional endothelial cells^1,13,66,67^. Although ETV2-directed reprogramming of iPSC-derived MPCs rapidly and efficiently yields ECs (this study and ref. ^1^), forced expression of ETV2 in fibroblasts was much less efficient^13,66,67^. Our atlas of efficient ETV2-directed MPC-to-EC specification will serve as a roadmap for future work to optimize EC differentiation and reprogramming strategies for therapeutic vascular regeneration.

## Data availability

The raw sequencing and related processed data have been deposited in GEO under the accession number (pending).

## Code availability

No custom code was used in this study. Analysis was done with publicly available pipelines using approaches described in the Methods.

## Acknowledgements

This work was supported by the National Institutes of Health (HL151450 to W.T.P. and J.M.M.) and American Heart Association (23POST1019621 to D.C.).

## Author contributions

D.C. and W.T.P. conceived and designed the experiments, interpreted results and wrote the manuscript with contributions from all of the authors. D.C. performed experiments, library preparation and bioinformatic analyses. X.F. assisted with experiments. K.W., L.G., and J.M.M. provided reagents, resources and interpreted results.

