## Supplemental data for "Pioneer factor ETV2 safeguards endothelial cell specification by recruiting the repressor REST to restrict alternative lineage commitment"

### Methods

All methods and tables will be available in the revised manuscript.

#### *Cell culture*

The human BJ273 iPSC line was sourced from Boston Children's Hospital's Stem Cell Core. TRE3G-ETV2 iPSCs were engineered from this parental line transfecting them with piggyBac expression plasmid (SBI #PB210PA-1) and the piggyBac transposon vector depicted in Fig. 1A. Stable cell lines were selected with puromycin (1 µg/ml), and clones were tested for pluripotency and Dox responsiveness.

iPSCs were maintained in mTeSR1 medium (StemCell Technologies, 85850) on Matrigel (Corning, 354277) pre-coated plates at 37 °C with 5% $CO_2$, and media was changed daily. iPSCs were passaged using TrypLE Select (Life Technologies, 12563011) and seeded with Rho-associated protein kinase (ROCK) inhibitor Y-27632 (StemCell Technologies, 72308) for the first day. Cells were frozen using mFreSR (StemCell Technologies, 05854) using a CoolCell LX (Corning, 432003) overnight at −80 °C and then in vapor-phase liquid nitrogen for long-term storage.

#### *iPSCs-derived endothelial cell differentiation*

iPSCs were differentiated into ECs using previously described differentiation medium[1]: Advanced DMEM/F12 (Thermo Fisher, 12634010), 1× GlutaMax supplement (Thermo Fisher, 35050061) and 60 µg/mL L-Ascorbic acid (Sigma-Aldrich, A8960). For differentiation, iPSCs were dissociated into single cells with TrypLE Select and plated on Matrigel pre-coated plates at a density of 60,000 iPSCs/$cm^2$ in mTeSR1 supplemented with 10 µM Y-27632. At Day 0, iPSCs were briefly washed with PBS and first differentiated into hMPCs using differentiation medium supplemented with CHIR99021(6 µM, StemCell Technologies, 100-1042) for 48 hours. At Day 2, hMPCs were dissociated and then seeded on a 60-mm Matrigel-coated dish in differentiation medium supplemented with VEGF-A (50 ng/mL, PeproTech, 100-20), fibroblast growth factor 2 (FGF-2, 50 ng/mL, PeproTech, 100-18B), EGF (10 ng/mL, PeproTech, AF-100-15), and SB431542 (10 µM, Selleckchem, S1067) for another 2 days. For Dox group, cells were treated with doxycycline (1 µg/mL, Sigma Aldrich, D3072) for 24 hours. Medium was changed every day throughout this protocol.

#### *RNA isolation and RT–qPCR analysis*

Total RNA was extracted using TRIzol reagent (Invitrogen, 15596026) and Zymo Research RNA Clean & Concentrator Kits (Zymo Research, ZR1013). Trizol aqueous phase was transferred to a Zymo-Spin IC Column and treated with DNase I. RNA was reverse transcribed to cDNA using SuperScript III First-Strand Synthesis SuperMix (Thermo Fisher, 11752050). RT-qPCR was carried out on a Bio-Rad CFX96 Touch Deep Well Real-Time PCR instrument using the SYBR Green PCR Master Mix (Thermo Fisher, 4368708). The primer sequences for qPCR analysis are listed in Supplementary Table 7. Relative gene expression was normalized to *GAPDH* and analyzed by $2^{-\Delta\Delta Ct}$ method.

#### *Flow cytometry*

Cells were dissociated using TrypLE Select, washed and resuspended in FACS buffer (PBS with 0.5% BSA). For surface antigens, live cells were stained with commercially available antibodies

(Supplementary Table 8) for 30 min on ice in the dark. Stained cells were washed twice with FACS buffer, filtered into a 35 µm strainer-capped tube (Falcon, 352235) and analysed by a BD LSRFortessa. FACS was performed on a BD FACSMelody Cell Sorter (BD Biosciences). Spectral overlap was compensated using UltraComp eBead Plus Compensation Beads (Life Technologies, 01-3333-42) with single fluorophore-conjugated antibodies. Single cells were gated using FSC-A and SSC-A. Flow cytometry data were analyzed using FlowJo software.

***Purification and expansion of ECs***

ECs obtained from differentiation of iPSCs were dissociated using TrypLE Select and sorted using PECAM1-conjugated magnetic microbeads (Miltenyi Biotec, 130-091-935) according to the manufacturer's instructions. The purified PECAM1-positive cells were seeded on 1% gelatin-coated plates and maintained in EGM2 medium (Promocell, C-22111) supplemented with 1× GlutaMax supplement and 10 µM SB431542. ECs were expanded for a maximum of 3 passages and used for functional assays.

***Immunofluorescent staining***

Immunofluorescent staining was performed as previously described[2]. Briefly, cells were washed with PBS followed by fixation with 4% paraformaldehyde for 15 min at room temperature. PBS containing 0.1% Triton X-100 was used for permeabilization. Then, cells were blocked for 1 h in blocking buffer (3% bovine serum albumin in PBS). After blocking, cells were incubated with primary antibodies (Supplementary Table 8) in blocking buffer at 4°C overnight. Cells were then washed three times followed by incubated with appropriate fluorescent secondary antibodies at RT for 2 h. Finally, cells were stained with a DAPI solution and mounted in ProLong Glass Antifade Mountant (Thermo Fisher, P36982). Images were taken using a Olympus FV3000R confocal microscope.

***Wound healing scratch assay***

Wound healing scratch assays were performed as described previously[3]. ECs were seeded into 6-well plates and grown to reach about 90% confluency on the day of the experiment. Monolayers were scratched using a pipette tip and washed three times with complete EGM2 medium. Scratches were imaged using a Keyence automated epifluorescent microscope equipped with a 10× objective. Cells were imaged after 8 and 16 h to determine their capacity to migrate in the cell free gap.

***Shear stress-induced polarization of iPSC-derived endothelial cells***

The resultant ECs were cultured in a 1% gelatin pre-coated 35-mm glass-bottom dishes (Nest) with EGM2 medium for 24 hours, either in static culture or a rotator. Confluent monolayers of ECs were subjected to orbital shear stress at a rotating frequency of 150 rpm using an orbital shaker positioned inside a cell culture incubator. After 24 hours of static or shear stress conditions, immunostaining was performed to quantify the degree of cell alignment. Only the cells in the periphery of the culture dish were imaged. After imaging, quantification of cell orientation angles was performed using ImageJ.

*NO production assay*

To determine the ability of the cells to produce nitric oxide (NO), cells were placed in fresh media containing 1 μM DAF-FM Diacetate (Thermo Fisher, D23844) or vehicle (DMSO). Cells were cultured for 30 min and then harvested for flow cytometric analysis. As a negative control, iPSCs were incubated in mTeSR1 containing DMSO or DAF-FM Diacetate. Geometric mean fluorescence intensities were calculated using FlowJo.

*Acetylated LDL (AcLDL) uptake assay*

To assess the ability of ECs to uptake AcLDL, ECs were incubated at 37°C for 4 hours with AcLDL conjugated to Alexa Fluor 488 (5 μg/mL, Thermo Fisher, L23380) in complete EGM2 medium. Following incubation, the ECs were washed with PBS to remove the excess stain and microscopic images were taken to evaluate the mean fluorescence in high-power fields. Representative image shown is selected among multiple independent experiments.

*Co-immunoprecipitation (Co-IP)*

Cells were lysed with Pierce IP Lysis Buffer (Thermo Scientific, 87787) supplemental with Halt Protease and Phosphatase Inhibitor Cocktail (Thermo Scientific, 78440). The lysates were centrifuged for 30 min at 16,000 g at 4°C, and the supernatants were transferred. 50 μL of lysate was used as the input. Dynabeads Protein G (Life Technologies, 10004D) were incubated with primary antibody (Supplementary Table 8) or IgG at 4℃ overnight. About 1 mg of cell lysate was added to the magnetic bead-antibody complex and incubated with rotation for 4°C overnight. The magnetic bead-antibody-antigen complex was pulled down and washed with PBS. The magnetic beads were resuspended in SDS-loading buffer and then heated up to 95 °C for 10 min, and proteins were analyzed by immunoblotting.

*Immunoblotting*

Total protein was extracted from cells using RIPA lysis buffer (Santa Cruz Biotechnology, SC-24948). The lysates were centrifuged for 30 min at 16,000 g at 4°C, and the supernatants were transferred. Protein concentration was determined using the Pierce BCA Protein Assay Kit (Life Technologies, 23227). Protein separation and immunoassay were conducted with the automated WES capillary based electrophoresis system (Wes; ProteinSimple). Target proteins were identified using primary antibodies (Supplementary Table 8) and immunoprobed using an HRP-conjugated secondary antibody (ProteinSimple). Chemiluminescent signals were detected and analyzed using Compass for SW (ProteinSimple) software.

*scRNA-seq data analysis*

*Data processing.*

FASTQ files were processed with the 10× Genomics CellRanger pipeline (v6.1.2). The raw reads were individually aligned to human reference genome GRCh38 (reference 2020-A) and quantified unique molecular identifiers (UMIs) counted by ‘cellranger count’ for each gene in each cell. The count matrix of all samples was imported into the R environment (v4.1) using the function ‘Read10X’ in Seurat R package (v4.1.0)[4]. We remove cells with <1200 genes, >8000 genes, or >10% mitochondrial reads. Only high-quality single-cell transcriptomes were used for subsequent analysis. The count matrix was classified into the Ctrl group (8 samples, 2 replicates each timepoint) and Dox group (8 samples, 2 replicates each timepoint) with a total 103,362 cells retained. A list of the number of cells in each sample is provided in Supplementary Table 1.

*Data integration and clustering.*

Standard Seurat data processing and normalization steps were performed for each group: SCTransform and RunPCA. A total of 3,000 genes were selected as integration features and used to identify 'integration anchors'. The data from both groups were integrated using IntegrateData functions in Seurat for batch correction. Then, we performed Seurat function RunPCA, FindNeighbors, FindClusters, and RunUMAP on the integrated data. UMAP embedding was computed for the integrated data using the top 40 principal components (PCs). Integration of transcriptomes from all timepoint or day 4 of Ctrl and Dox was performed. DoubletFinder[5] was used with default parameters to remove potential doublets. The overlapping embedding of Ctrl and Dox on Day 2 suggested effective removal of batch effects. Expression of marker genes was denoted by different color shades and superimposed over the entire cell population, as visualized on a UMAP projection. Gene-set signature scores were computed using the AddModuleScore function in Seurat.

*Differential gene expression tests.*

Each cluster's differentially expressed genes (DEGs) were identified using the FindAllMarkers function with default parameters (min.pct set to 0.1 and logfc.threshold set to 0.25). Top 5 DEGs with a $|\log_2$ fold change$| > 1.5$ from the Wilcoxon rank-sum test were displayed using the Dotplot function. For pairwise comparisons between Ctrl and Dox cells on Day 3, DEGs were identified using the FindMarkers functions. DEGs with $|\log_2$ fold change$| > 1$ and Padj $< 0.05$ were shown in the scatter plot. All heat maps were generated using Seurat's DoHeatmap function.

*Ontology annotation.*

We performed gene ontology (GO) analysis on lists of differentially expressed genes using ClusterProfiler package (v4.2.2)[6]. The Simplify function was used to remove redundant GO terms. GO terms with q value less than 0.05 were defined as significantly enriched.

***scATAC-seq data analysis***

*Data processing and clustering.*

FASTQ files were processed with the 10× Genomics CellRanger ATAC pipeline (v2.1.0). Raw reads were individually aligned to the human reference genome GRCh38 (reference 2020-A). The command 'cellranger-atac count' was used to obtain the fragment file for each sample. Downstream analysis was performed in R (v4.2.0), Seurat (v4.1.0), Signac (1.9.0)[7] and ArchR (1.0.2)[8]. The union peak list was generated using all peak lists. The barcoded fragments files were used as input data for analysis using 'FeatureMatrix' and 'CreateChromatinAssay' functions in Signac. The different samples were pooled together. Quality-control metrics were calculated and cells were filtered as follows: number of fragments, 1,000 - 30,000; blacklist_fraction < 0.05; nucleosome_signal < 4; and TSS enrichment score > 2. In addition to Signac metrics, ArchR was also used to perform quality control and identify doublets. The matrix was classified into Ctrl (6 samples) and Dox (6 samples) with a total 67,706 cells retained. A list of the number of cells in each sample is provided in Supplementary Table 1. Standard Iterative Latent Semantic Indexing (LSI) dimension reduction and clustering was performed as follows using default parameters: RunTFIDF, FindTopFeatures, RunSVD, RunUMAP, FindNeighbors and FindClusters. UMAP was used to visualize cell embedding for all cells.

*Integrative analysis of scATAC-seq and scRNA-seq.*

Seurat's canonical correlation analysis (CCA) was used to align scRNA-seq and scATAC-seq datasets by comparing scATAC-seq gene activity with the scRNA-seq gene expression matrix. For this purpose, gene activity in the cells profiled by scATAC-seq was calculated using the 'GeneActivity' function in Signac followed by Normalize Data with the LogNormalize method. The highly variable genes from the scRNA-seq dataset were used as input to 'FindTransferAnchors' function in Seurat with reduction method 'cca' to identify integration anchors. We then assigned each of the scATAC-seq cells a cell type subcluster identity from the matching scRNA-seq data and an associated label prediction score using the 'TransferData' functions in Seurat with default parameters. To ensure the prediction accuracy, only the cells with a prediction score higher than 0.5 were retained for further analysis. Finally, the annotation of scATAC-seq clusters was defined based on the identity of its cells.

Downstream analysis was performed using the ArchR package, unless otherwise noted. The GeneScoreMatrix and GeneIntegrationMatrix in scATAC-seq was created after the Seurat label transfer procedure. To identify the marker genes in scATAC-seq, gene expression in each cell type was calculated using Wilcoxon test with $| \log_2$ fold change $| > 1$ and false discovery rate (FDR) $< 0.05$. Peak-to-gene correlation analysis for the feature genes was computed to identify putative regulatory relationships by correlating peak accessibility to GeneIntegrationMatrix across scATAC-seq metacells. This procedure was invoked using the 'addPeak2GeneLinks' function. Only links with correlation > 0.45 and FDR < 1e4 were visualized (Supplementary Table 3). Genome coverage tracks displaying the pseudo-bulk coverage patterns were generated using 'plotBrowserTrack' function.

*Identification of differentially accessible peaks*

scATAC-seq data were used to generate pseudo-bulk replicates based on cluster identities. Pseudo-bulk peak calling was performed within each cluster using MACS2[9] using the 'addReproduciblePeakSet'. ArchR's default iterative overlap peak merging procedure was then applied to merge all peak calls and generate a reproducible peak set. Differentially accessible marker peaks in each cluster were identified by comparing each cluster cell using Wilcoxon test with $| \log_2$ fold change $| > 1$ and FDR $< 0.01$. For pairwise comparisons, the differentially accessible peaks were calculated using the Wilcoxon test with the same threshold. Gene Ontology analysis of regions was performed using GREAT[10] with its basal plus extension option and default parameters.

**Statistics**

Statistical analyses were performed in R or Prism 10 (GraphPad). Results are presented as the mean ± SEM unless noted. Statistical tests are indicated in the figure legends.

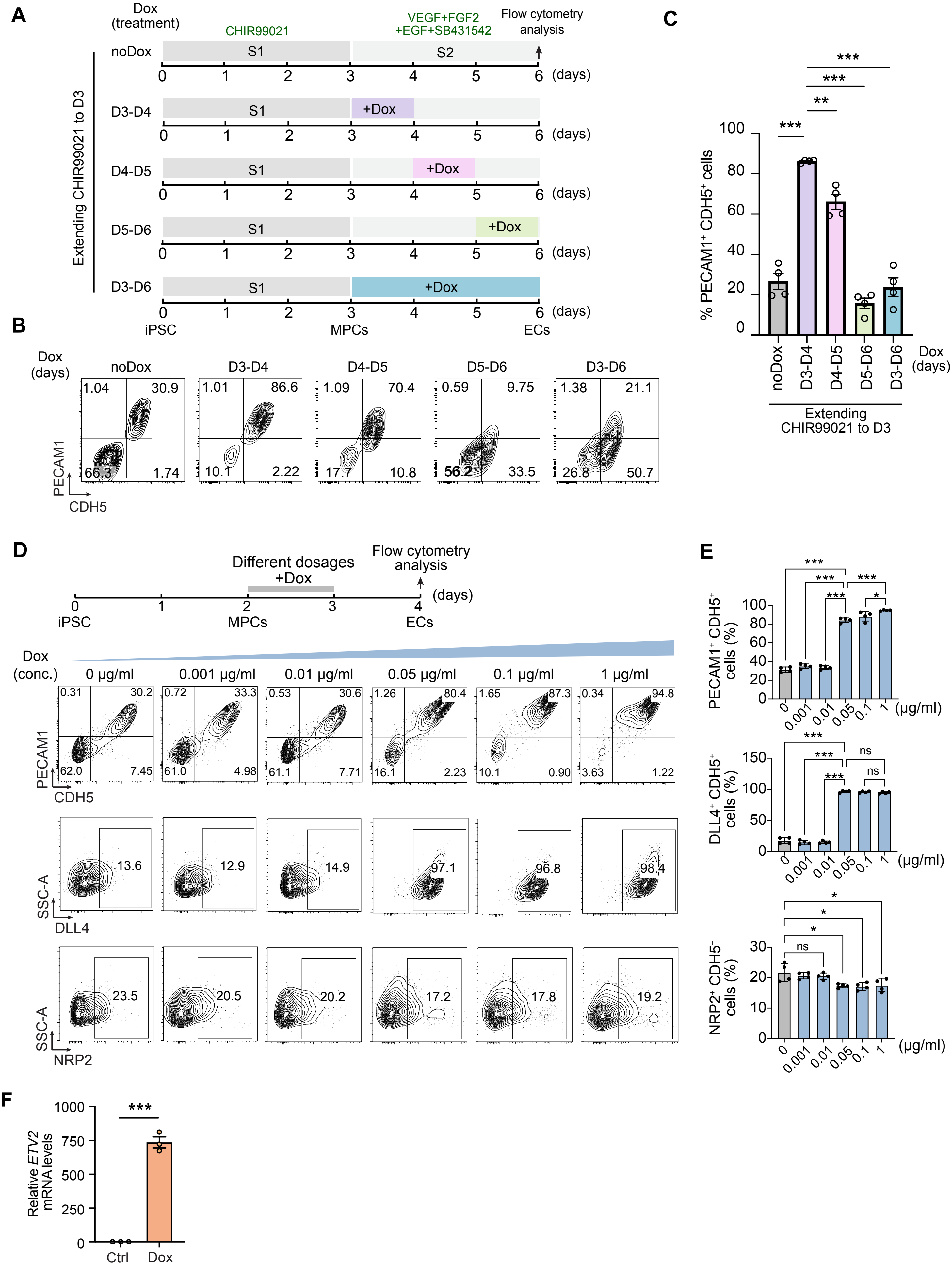

A
Dox (treatment)
CHIR99021
VEGF+FGF2 +EGF+SB431542
Flow cytometry analysis
noDox
D3-D4
D4-D5
D5-D6
D3-D6
S1
S2
+Dox
(days)
Extending CHIR99021 to D3
iPSC
MPCs
ECs
B
Dox (days)
noDox
D3-D4
D4-D5
D5-D6
D3-D6
1.04
30.9
66.3
1.74
1.01
86.6
10.1
2.22
1.09
70.4
17.7
10.8
0.59
9.75
56.2
33.5
1.38
21.1
26.8
50.7
PECAM1
CDH5
C
% PECAM1⁺ CDH5⁺ cells
***
**
Dox (days)
Extending CHIR99021 to D3
D
Different dosages +Dox
Flow cytometry analysis
(days)
iPSC
MPCs
ECs
Dox (conc.)
0 μg/ml
0.001 μg/ml
0.01 μg/ml
0.05 μg/ml
0.1 μg/ml
1 μg/ml
0.31
30.2
62.0
7.45
0.72
33.3
61.0
4.98
0.53
30.6
61.1
7.71
1.26
80.4
16.1
2.23
1.65
87.3
10.1
0.90
0.34
94.8
3.63
1.22
PECAM1
CDH5
13.6
12.9
14.9
97.1
96.8
98.4
SSC-A
DLL4
23.5
20.5
20.2
17.2
17.8
19.2
SSC-A
NRP2
E
PECAM1⁺ CDH5⁺ cells (%)
DLL4⁺ CDH5⁺ cells (%)
NRP2⁺ CDH5⁺ cells (%)
(μg/ml)
ns
*
F
Relative ETV2 mRNA levels
Ctrl
Dox

**Extended Data Fig. 1. ETV2 is required during EC formation. (A)** Scheme of 6 day EC differentiation protocol showing timing of Dox addition (1 µg/ml). Cells were treated without Dox, or treated with Dox for indicated length of time. **(B,C)** Flow cytometry analysis (B) and quantification (C) for PECAM1 and CDH5 expression at Day 6 of differentiation. One-way ANOVA with Tukey's multiple comparison test. (n = 4). **(D,E)** Scheme of 4 day EC differentiation protocol using different Dox dosages. Cells treated with Dox at Day 2 for 24 hours were analyzed for flow cytometry of PECAM1$^+$ CDH5$^+$ ECs, DLL4$^+$ CDH5$^+$ artery ECs, or NRP2$^+$ CDH5$^+$ venous ECs at Day 4. One-way ANOVA with Tukey’s multiple comparison test. (n = 4). **(F)** Real-time PCR analysis of *ETV2* mRNA expression in each group on day 3. Two-tailed unpaired Student’s t-test. (n = 3). Data are mean ± s.e.m. *, P<0.05; **, P<0.01; ***, P<0.001, ns, not significant.

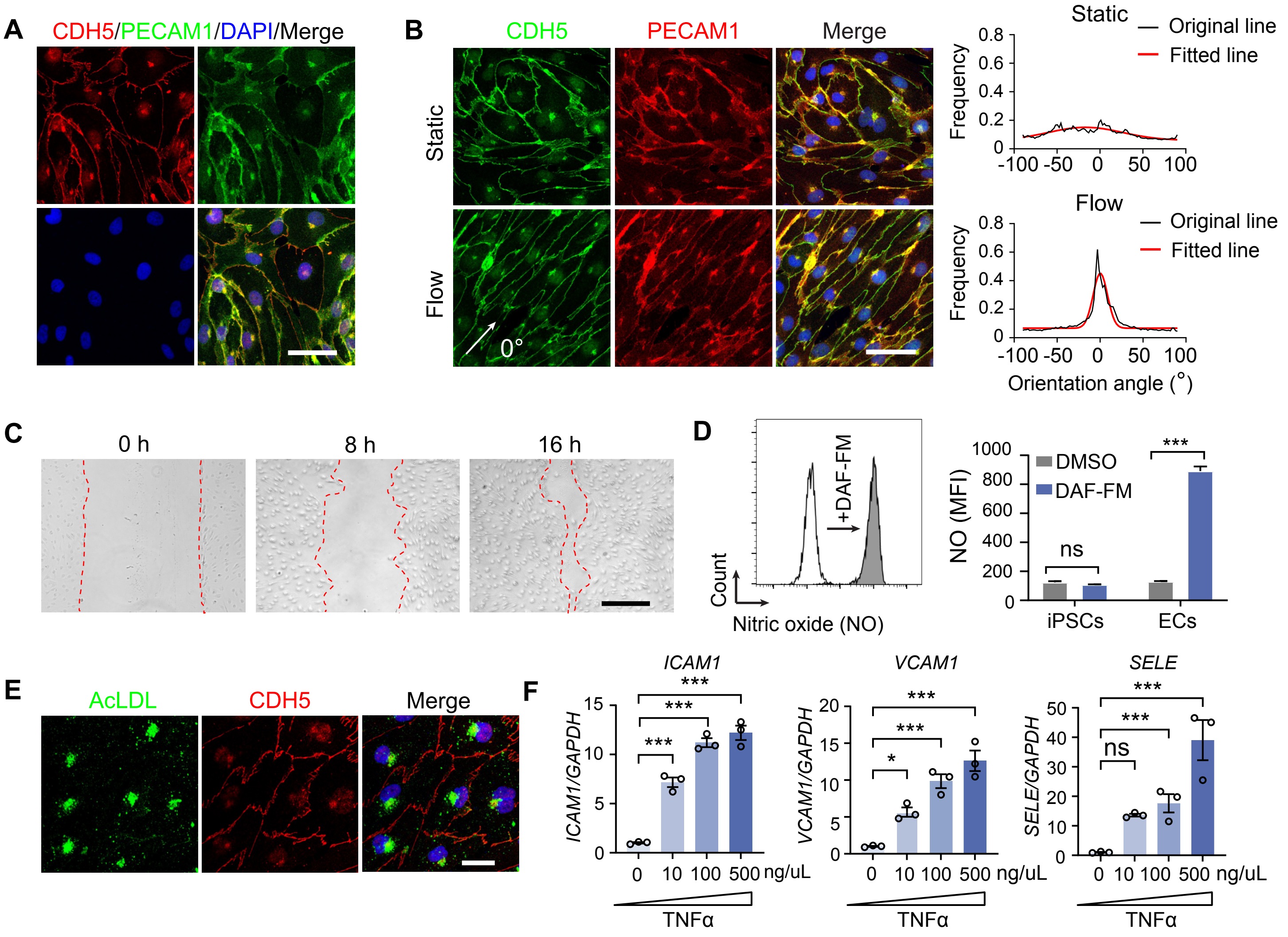


**Extended Data Fig. 2. Functional characterization of cultured iPSC-ECs with ETV2 overexpression. (A)** iPSC-ECs immunostained for CDH5 (red), PECAM1 (green), and DAPI (blue). Scale bar, 50 µm. **(B)** Alignment of iPSC-ECs under static and flow conditions. 0° represents the direction of flow. Scale bars, 50 µm. **(C)** Migration of iPSC-ECs in the wound scratch assay. Scale bar, 300 µm. **(D)** NO production of iPSC-ECs measured by flow cytometry upon exposure to DAF-FM. MFI, mean fluorescence intensity. Two-way ANOVA (n = 3). **(E)** Acetylated LDL (AcLDL, green) uptake by iPSC-ECs. Scale bar, 25 µm. **(F)** Expression of cell surface adhesion molecules *ICAM1*, *VCAM1*, and *E-Selectin (SELE)* in iPSC-ECs when stimulated with 10, 100, or 500 ng/uL tumor necrosis factor-alpha (TNFα). One-way ANOVA with Dunnett's multiple comparison test. (n = 3). Data are mean ± s.e.m. *, P<0.05; ***, P<0.001; ns, not significant.

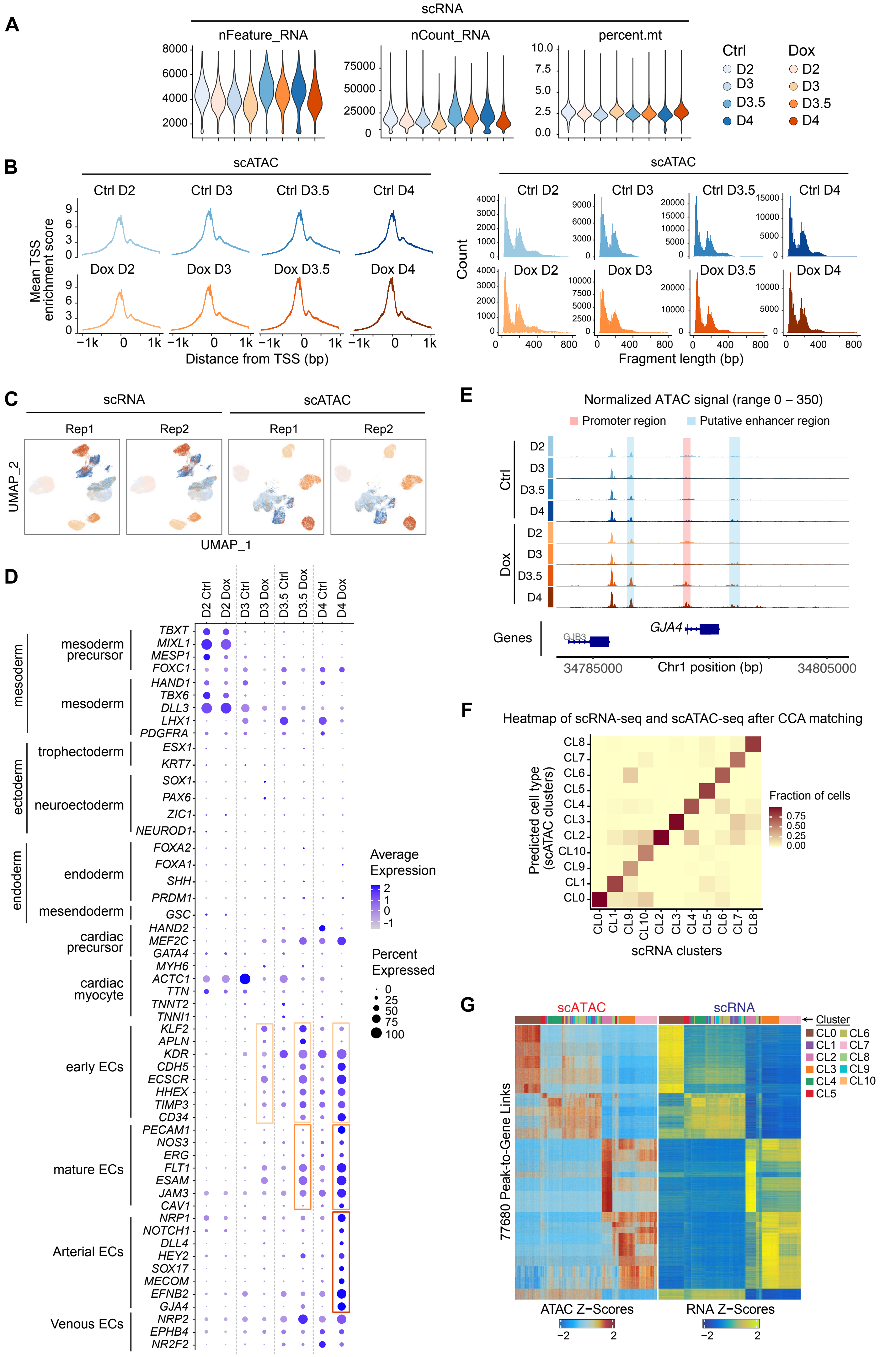

A
scRNA
nFeature_RNA
nCount_RNA
percent.mt
Ctrl
Dox
D2
D3
D3.5
D4
B
scATAC
Mean TSS enrichment score
Ctrl D2
Ctrl D3
Ctrl D3.5
Ctrl D4
Dox D2
Dox D3
Dox D3.5
Dox D4
Distance from TSS (bp)
Count
Fragment length (bp)
C
scRNA
scATAC
Rep1
Rep2
UMAP_2
UMAP_1
D
mesoderm
mesoderm precursor
trophectoderm
ectoderm
neuroectoderm
endoderm
mesendoderm
cardiac precursor
cardiac myocyte
early ECs
mature ECs
Arterial ECs
Venous ECs
Average Expression
Percent Expressed
E
Normalized ATAC signal (range 0 – 350)
Promoter region
Putative enhancer region
Genes
GJB3
GJA4
34785000
Chr1 position (bp)
34805000
F
Heatmap of scRNA-seq and scATAC-seq after CCA matching
Predicted cell type (scATAC clusters)
Fraction of cells
scRNA clusters
G
scATAC
scRNA
Cluster
77680 Peak-to-Gene Links
ATAC Z−Scores
RNA Z−Scores

**Extended Data Fig. 3. Quality control of scRNA-seq and scATAC-seq data and gene expression of representative cell type markers. (A)** Quality control metrics of scRNA seq data for cell samples from Ctrl and Dox group. Number of genes, number of unique molecular identifiers (UMIs), and mitochondrial gene percentage is plotted for each sample. **(B)** Quality control metrics of scATAC-seq data for cell samples from Ctrl and Dox groups. Left, Aggregate ATAC signal at all TSS regions for the cells passing QC thresholds for each sample. Right, fragment length distribution for each sample. **(C)** Biological replicates of scRNA-seq and scATAC-seq data from D2, D3, D3.5 and D4 projected onto UMAP space. Cells are colored by sample collection time.**(D)** Dot plots of quantitative bulk changes in gene expression between Ctrl and Dox cells for markers of mesoderm, ectoderm, endoderm, cardiac precursors, early ECs, mature ECs, and genes involved in artery EC and venous EC development. **(E)** Genome tracks of time point-resolved aggregate scATAC-seq data around the *GJA4* gene loci. **(F)** Heatmap showing the cluster-cluster mapping between scRNA-seq and scATAC-seq clusters after canonical correlation analysis (CCA) matching. **(G)** Heatmap of statistically significant distal 77,680 peak-to-gene links identified across the dataset with ArchR (see Methods). Each row represents expression of a gene (scRNA, right) correlated to accessibility of a distal peak (scATAC, left).

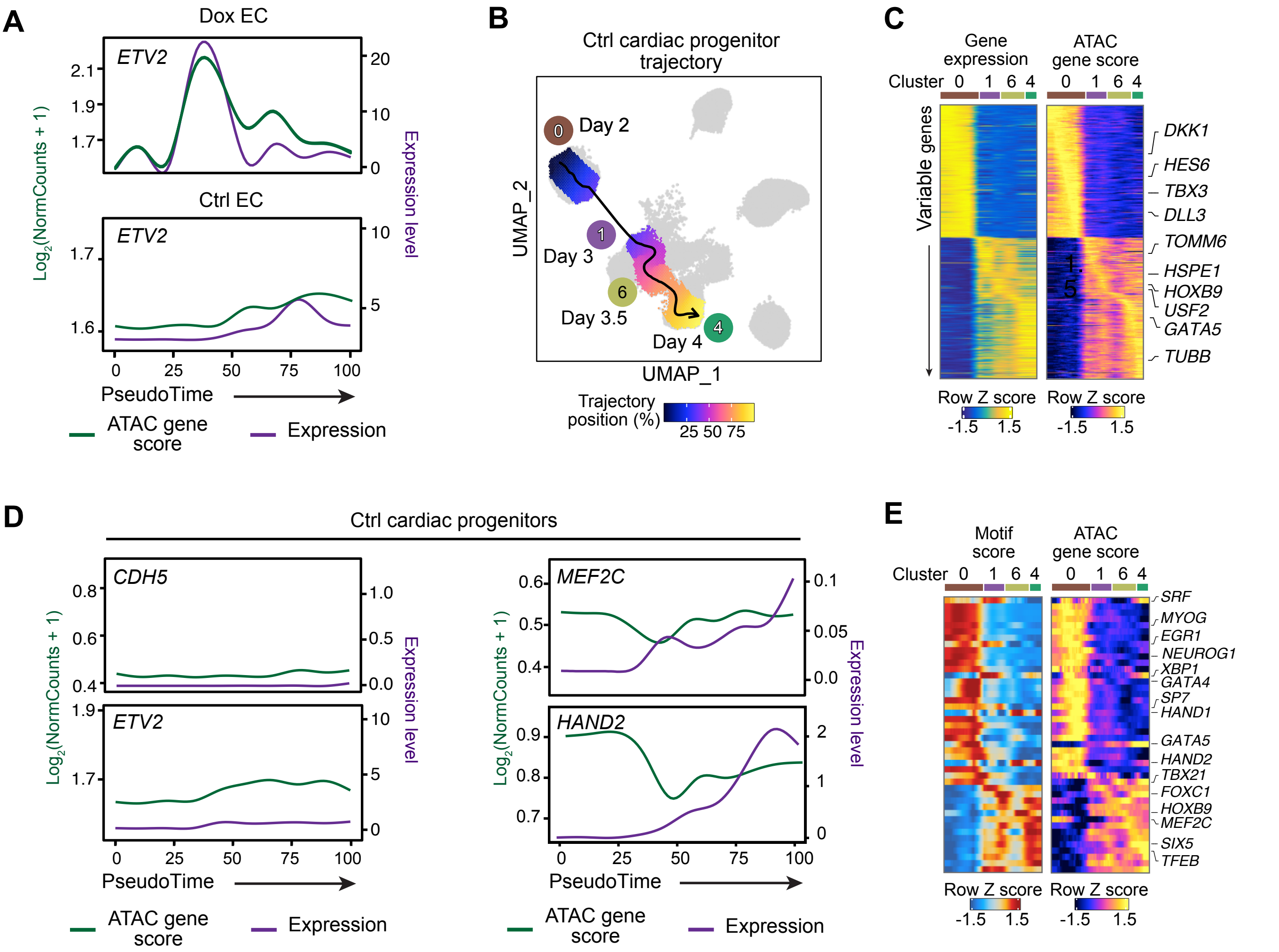


**Extended Data Fig. 4. Ctrl trajectory terminating on cardiac progenitor cells, related to Fig. 3.** **(A)** Gene expression and chromatin-derived gene accessibility score (ATAC gene score) dynamics of *ETV2* in Dox EC and Ctrl EC trajectory across pseudotime. **(B)** UMAP visualization of the cardiac progenitor trajectory of Ctrl cells. Each cell is colored by its pseudotime and each cluster is annotated by sample collection time. The smoothed arrow represents a visualization of the interpreted trajectory in the UMAP embedding. **(C)** Heatmap of scRNA-seq gene expression (left) and scATAC-seq gene score (right) along the cardiac progenitor trajectory. Genes displayed showed the greatest variation along the trajectory. Cell clusters are labeled according to their position along the pseudotime. **(D)** Gene expression and chromatin-derived gene accessibility score (ATAC gene score) dynamics of the *CDH5*, *ETV2*, and cardiac progentior genes *MEF2C* and *HAND2*. **(E)** Heatmap of TF regulators for which ATAC gene score is positively correlated with chromVAR TF deviation (motif score) across the Ctrl CP trajectory, ordered by pseudotime.

**A**

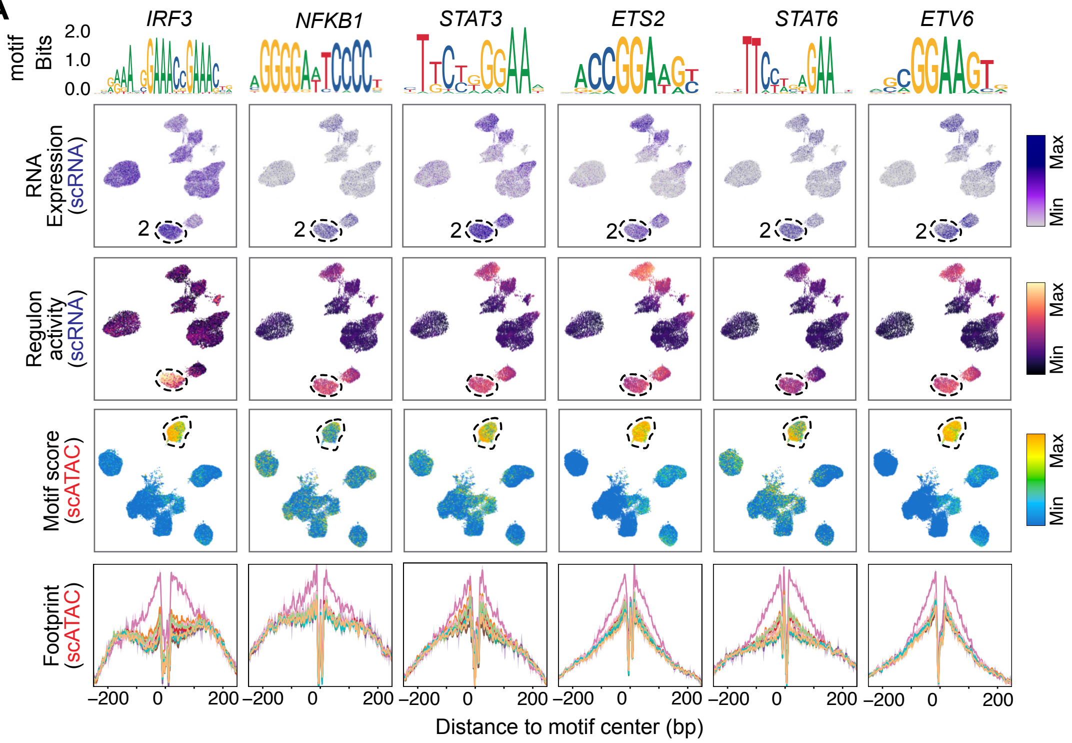


**B**

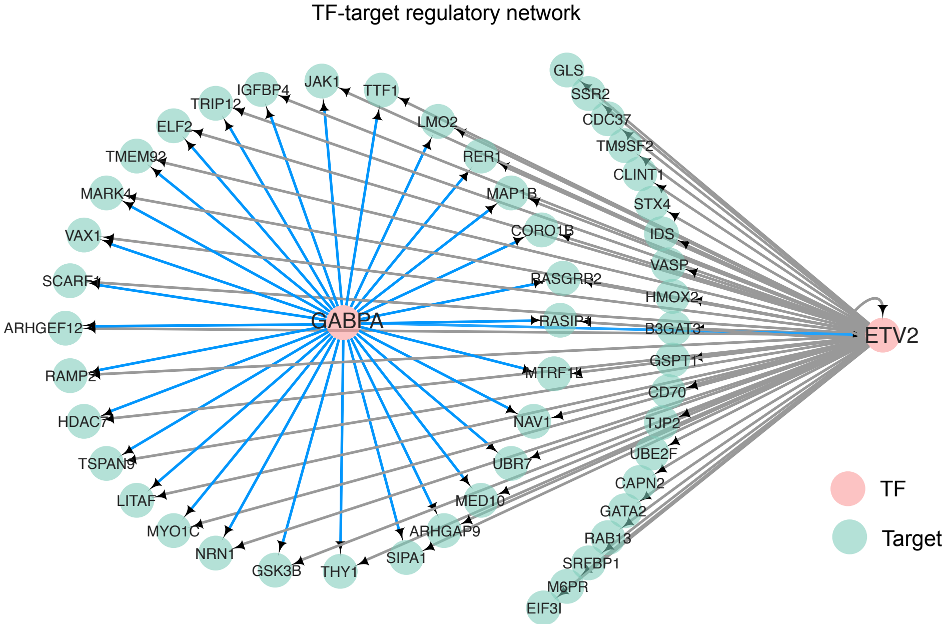


**Extended Data Fig. 5. ETV2 drives endothelial specification in collaboration with other TFs in endothelial progenitors. (A)** TFs with enriched motif scores or regulon activity in CL2, ETV2$^{OE}$ EP-1. 1st row: feature plots of scRNA-seq gene expression; 2nd row: regulon activity; 3rd row: scATAC-seq based chromVAR TF deviation (motif score); 4th row: Tn5 bias-adjusted TF footprints. Gene expression was positively correlated with motif score across differentiation, representing positive TF regulators of differentiation. **(B)** Visualization of TF-target regulatory networks formed by *ETV2* and *GABPA*. The pink nodes represent the TFs. The light green nodes represent the targets. The arrows, representing the connections between each of the TFs and their predicted target genes (GABPA, blue line; ETV2, grey line).

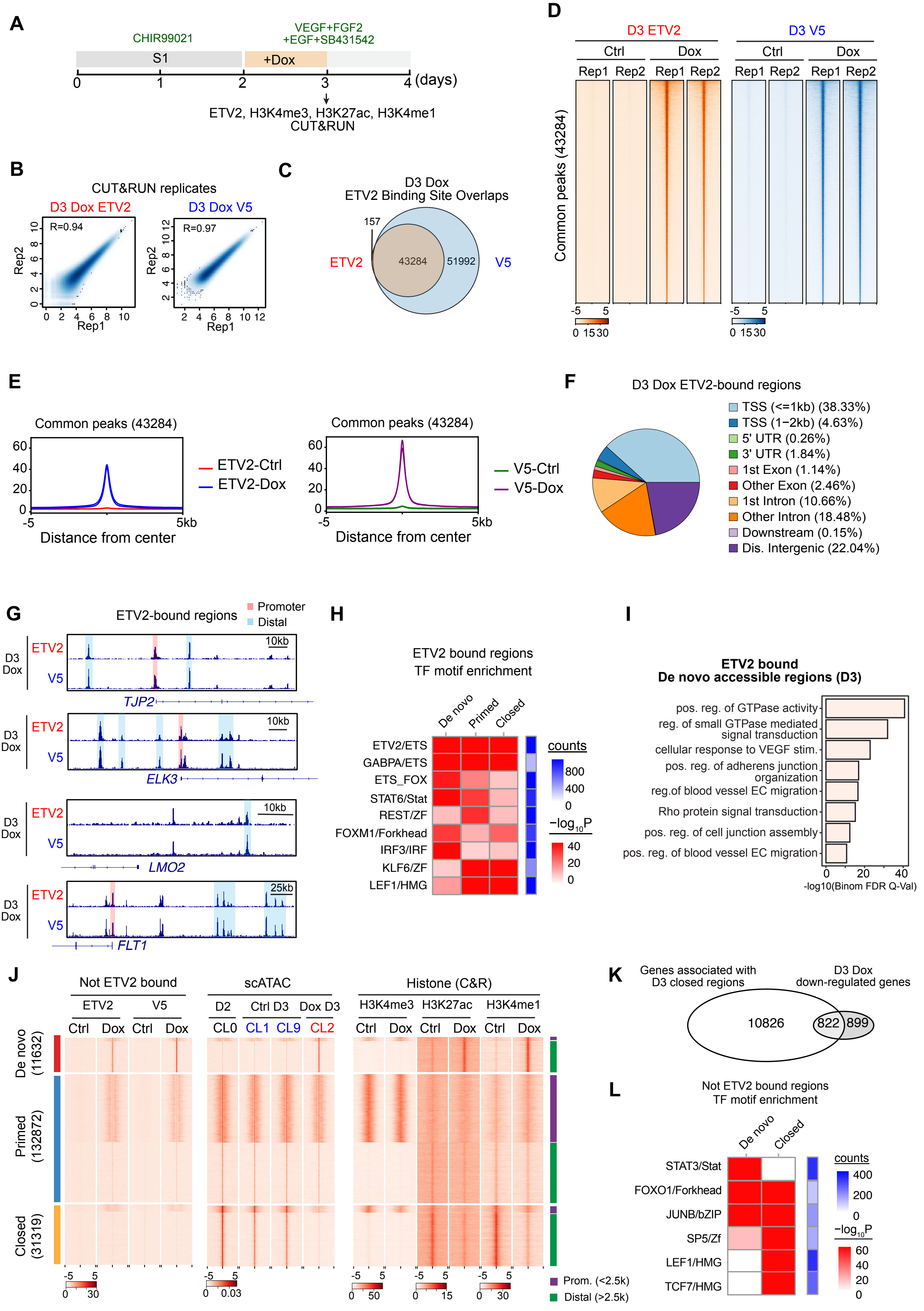
A
CHIR99021
VEGF+FGF2
+EGF+SB431542
S1
+Dox
0 1 2 3 4 (days)
ETV2, H3K4me3, H3K27ac, H3K4me1
CUT&RUN
B
CUT&RUN replicates
D3 Dox ETV2
D3 Dox V5
R=0.94
R=0.97
Rep2
Rep1
C
D3 Dox
ETV2 Binding Site Overlaps
157
ETV2
43284
51992
V5
D
D3 ETV2
D3 V5
Ctrl
Dox
Rep1 Rep2
Common peaks (43284)
E
Common peaks (43284)
ETV2-Ctrl
ETV2-Dox
V5-Ctrl
V5-Dox
Distance from center
F
D3 Dox ETV2-bound regions
TSS (<=1kb) (38.33%)
TSS (1−2kb) (4.63%)
5' UTR (0.26%)
3' UTR (1.84%)
1st Exon (1.14%)
Other Exon (2.46%)
1st Intron (10.66%)
Other Intron (18.48%)
Downstream (0.15%)
Dis. Intergenic (22.04%)
G
ETV2-bound regions
Promoter
Distal
D3 Dox
ETV2
V5
10kb
25kb
TJP2
ELK3
LMO2
FLT1
H
ETV2 bound regions
TF motif enrichment
De novo
Primed
Closed
ETV2/ETS
GABPA/ETS
ETS_FOX
STAT6/Stat
REST/ZF
FOXM1/Forkhead
IRF3/IRF
KLF6/ZF
LEF1/HMG
counts
800
400
0
−log10P
40
20
0
I
ETV2 bound
De novo accessible regions (D3)
pos. reg. of GTPase activity
reg. of small GTPase mediated signal transduction
cellular response to VEGF stim.
pos. reg. of adherens junction organization
reg.of blood vessel EC migration
Rho protein signal transduction
pos. reg. of cell junction assembly
pos. reg. of blood vessel EC migration
0 20 40
-log10(Binom FDR Q-Val)
J
Not ETV2 bound
scATAC
Histone (C&R)
ETV2
V5
Ctrl
Dox
D2
Ctrl D3
Dox D3
CL0
CL1
CL9
CL2
H3K4me3
H3K27ac
H3K4me1
De novo (11632)
Primed (132872)
Closed (31319)
Prom. (<2.5k)
Distal (>2.5k)
K
Genes associated with D3 closed regions
D3 Dox down-regulated genes
10826
822
899
L
Not ETV2 bound regions
TF motif enrichment
De novo
Closed
STAT3/Stat
FOXO1/Forkhead
JUNB/bZIP
SP5/Zf
LEF1/HMG
TCF7/HMG
counts
400
200
0
−log10P
60
40
20
0

**Extended Data Fig. 6. ETV2 directly binds promoters and enhancers of target genes.** **(A)** Timing of the ETV2, H3K4me3, H3K27ac and H3K4me1 CUT&RUN analyses. ETV2 tagged with V5 was induced at day 2 by addition of Dox, and CUT&RUN was performed at day 3. **(B)** Scatter plots comparing the Dox ETV2 and V5 CUT&RUN signals at day 3 during EC differentiation between two biological replicates. Pearson correlation coefficients are shown. **(C)** Venn diagram showing overlap between Dox D3 ETV2 and V5 CUT&RUN peaks. Overlapping peaks were defined as high confidence ETV2 regions. **(D)** ETV2 (orange) and V5 (blue) CUT&RUN signal at high confidence ETV2 regions. Signal was restricted to Dox cells. Each row represents a peak region (peak center ± 5 kb). **(E)** Aggregation plots of D3 ETV2 and V5 CUT&RUN signal at high confidence ETV2 regions in Ctrl and Dox cells. **(F)** Genomic distribution of high confidence ETV2 regions in Dox cells at day 3. The number in brackets is the percentage of ETV2 peaks in each region. **(G)** Genome tracks of Dox D3 ETV2 CUT&RUN data around the *TJP2*, *ELK3*, *LMO2*, and *FLT1* gene loci. **(H)** Motifs enriched in ETV2-bound de novo, primed, and closed regions (presented in Fig. 5D). **(I)** GREAT analysis of GO biological process terms significantly enriched at ETV2-bound de novo accessible regions (presented in Fig. 5D). **(J)** Heatmap representing the signal for indicated chromatin features at D3 Dox de novo, primed, and closed regions that were not occupied by ETV2. **(K)** Venn diagrams showing the overlap of genes associated with closed regions and D3 Dox down-regulated genes. **(L)** Motifs enriched in D3 Dox de novo accessible or closed regions that were not bound by ETV2.

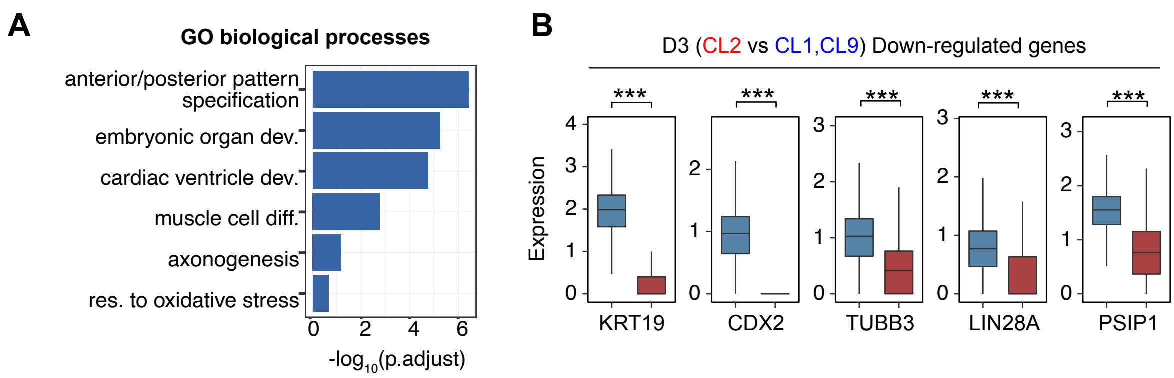


**Extended Data Fig. 7. Downregulated genes in ETV2$^{OE}$ endothelial progenitor cells. (A)** GO term analysis was performed on D3 Dox downregulated genes compared to D3 Ctrl. GO categories are ordered on the basis of -log$_{10}$(p.adj) value. **(B)** Expression of representative downregulated genes in D3 Ctrl and Dox scRNA data.Two way Student's t-test. ***, P<0.001.

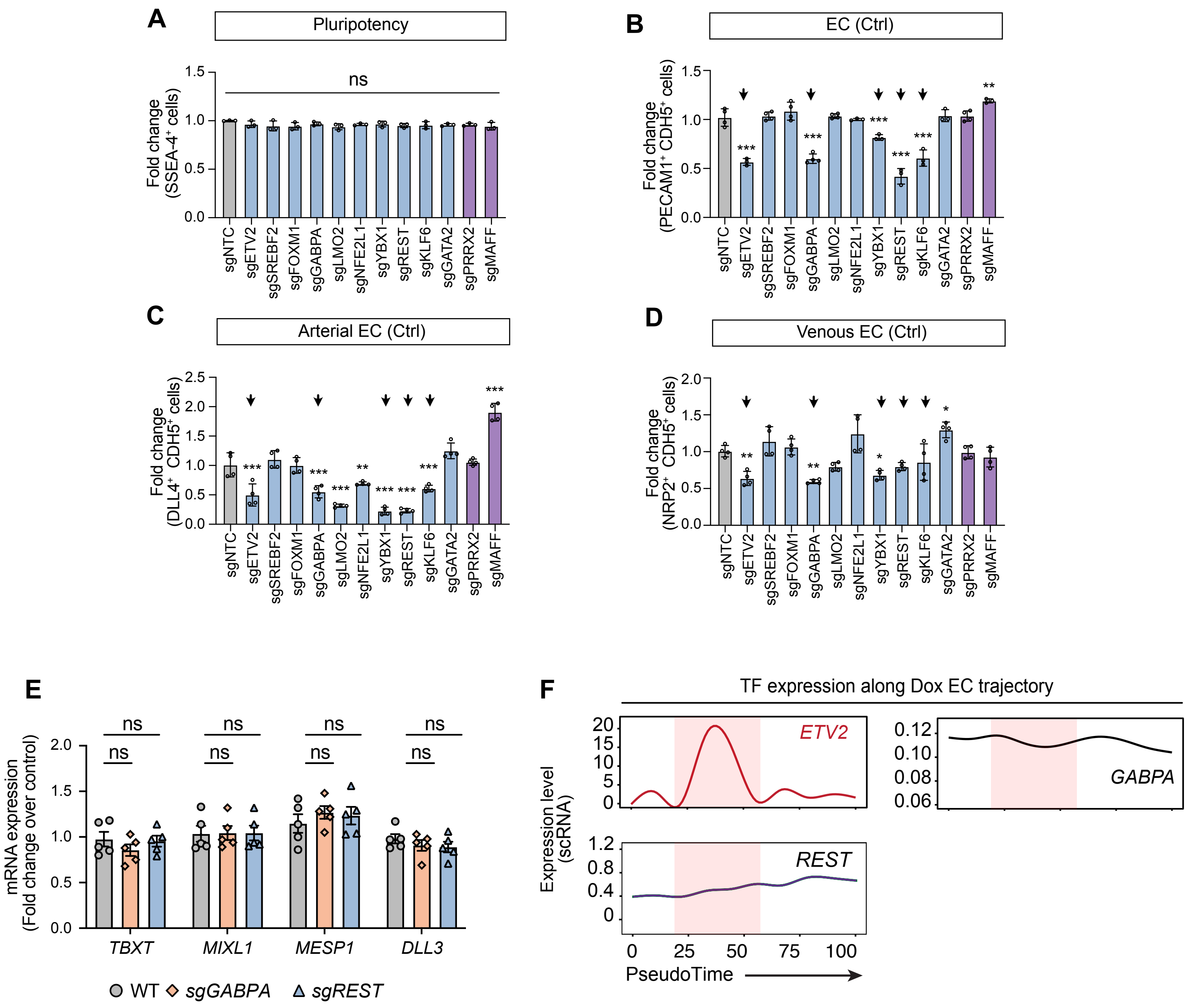
A
Pluripotency
ns
Fold change (SSEA-4+ cells)
B
EC (Ctrl)
Fold change (PECAM1+ CDH5+ cells)
C
Arterial EC (Ctrl)
Fold change (DLL4+ CDH5+ cells)
D
Venous EC (Ctrl)
Fold change (NRP2+ CDH5+ cells)
sgNTC
sgETV2
sgSREBF2
sgFOXM1
sgGABPA
sgLMO2
sgNFE2L1
sgYBX1
sgREST
sgKLF6
sgGATA2
sgPRRX2
sgMAFF
E
mRNA expression (Fold change over control)
TBXT
MIXL1
MESP1
DLL3
WT
sgGABPA
sgREST
F
TF expression along Dox EC trajectory
ETV2
GABPA
REST
Expression level (scRNA)
PseudoTime

**Extended Data Fig. 8. Validation of top candidates in EC differentiation. (A)** Flow cytometry for $SSEA4^+$ in iPSCs that were transfected with sgNTC, or transfected with sgRNAs targeting to the indicated gene. One-way ANOVA with Dunnett's multiple comparison test to sgNTC control. n = 3. **(B-D)** Validation of screening hits. iPSCs were transfected with a non-targeting control sgRNA (sgNTC) or sgRNAs targeting to the indicated gene. Differentiation efficiency to $PECAM1^+$ $CDH5^+$ ECs (B), $DLL4^+$ $CDH5^+$ aECs (C), or $NRP2^+$ $CDH5^+$ vECs (D) under Ctrl conditions was measured by flow cytometry. n = 3 or 4. **(E)** RT-qPCR analysis of *TBXT*, *MIXL1*, *MESP1* and *DLL3* mRNA expression in each group on day 2. Two-way ANOVA. n = 5. **(F)** RNA expression dynamics of *ETV2*, *GABPA* and *REST* across the Dox EC trajectory (presented in Fig. 3E). The red shaded area indicates the ETV2 expression window. Data are mean ± s.e.m. *, P<0.05. **, P<0.01. ***, P<0.001. ns, not significant.

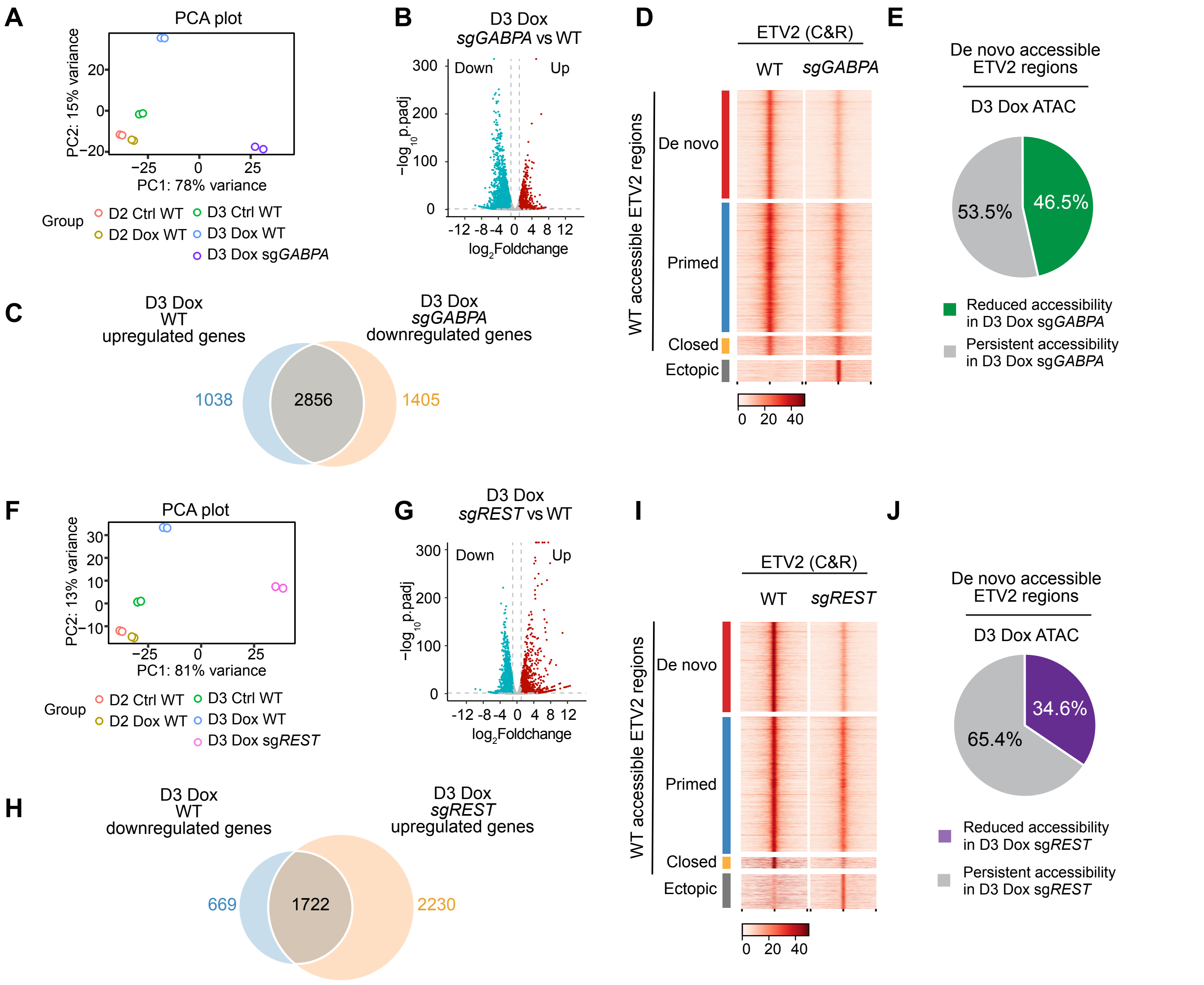


**Extended Data Fig. 9. GABPA and REST are required for chromatin opening and transcription activity. (A)** PCA plot summarizing RNA-seq of WT and *GABPA* knockout cells on D2 or D3. **(B)** Volcano plot showing DEGs identified in Dox D3 *GABPA* knockout cells versus Dox D3 WT cells. Genes with | $Log_2$Foldchange | > 1 and $P_{adj} < 0.05$ are considered DEGs. Significantly upregulated or downregulated genes are colored red or teal, respectively. **(C)** Overlap between genes up-regulated in D3 Dox (vs. Ctrl) and down-regulated in Dox D3 *GABPA* knockout (vs. WT). **(D)** ETV2 binding signals in Dox D3 WT and Dox D3 *GABPA* knockout. Regions are group by chromatin occupancy class in Dox D3 WT (de novo, primed, closed) and ectopic binding sites in *GABPA* knockout. **(E)** Effect of *GABPA* knockout on chromatin accessibility. Most ETV2-bound de novo accessible regions lost accessibility in Dox D3 *GABPA* knockout. **(F)** PCA plot summarizing RNA-seq of WT and *REST* knockout cells on D2 or D3. **(G)** Volcano plot showing DEGs identified in Dox D3 *REST* knockout cells versus Dox D3 WT cells. Genes with | $Log_2$Foldchange | > 1 and $P_{adj} < 0.05$ are considered DEGs. Significantly upregulated or downregulated genes are colored red or teal, respectively. **(H)** Overlap between genes down-regulated in D3 Dox (vs. Ctrl) and up-regulated in Dox D3 *REST* knockout (vs. WT). **(I)** ETV2 binding signals in Dox D3 WT and Dox D3 *REST* knockout cells. Regions are group by chromatin occupancy class in Dox D3 WT (de novo, primed, closed) and ectopic binding sites in *REST* knockout. (**J**) Effect of *REST* knockout on chromatin accessibility. Many ETV2-bound de novo accesible regions lost accessibility in Dox D3 *REST* knockout.
